# Sustained mechanical tension governs fibrogenic activation of tendon stromal cells in systemic sclerosis

**DOI:** 10.1101/2021.06.11.445955

**Authors:** Amro A. Hussien, Robert Knell, Florian Renoux, Stefania L. Wunderli, Barbara Niederoest, Jasper Foolen, Oliver Distler, Jess G. Snedeker

## Abstract

Fibrosis is a pathological outcome of aberrant repair responses in systemic sclerosis and affects many tissues, including tendons. Progressive matrix stiffening is a key feature of this pathological remodeling. How dysregulated tissue mechanics contribute to the persistence of the fibrotic phenotype has been obscured by limited availability of experimental tissue models that are both controllable and capture essential aspects of the tendon biophysical niche. Here, we developed a modular, cantilever-based platform that allows culture of 3D tendon-like constructs under easily variable static tension, emulating this central tendon-specific structure function relationship. The system reveals that elevated matrix tension instigates fibroblast-to-myofibroblast activation eliciting scar-like phenotypes *in vitro*. By using this mechano-culture system and preclinical and clinical models of systemic sclerosis, we further show that 3D matrix stiffness is inversely correlated with the transcription of major pro-fibrotic collagens, but positively correlate with the expression of markers of stromal-immune interactions. Co-culture of tendon stromal fibroblasts and bone marrow-derived macrophages override stiffness-mediated downregulation of matrix transcription, suggesting that normal tension mediated checkpoints are superseded by the local tissue immune state. Our study highlights the power of 3D reductionist approaches in dissecting the contribution of the elevated matrix tension to the positive feedforward loops between activated fibroblasts and progressive ECM stiffening in systemic sclerosis.

## Introduction

Systemic sclerosis (SSc) is a chronic immune-mediated rheumatic disease, characterized by microvascular damage, non-resolving inflammation, and progressive fibrosis of connective tissues. (1–4) The disease represents significant clinical and socioeconomic burdens to patients, their caretakers and healthcare systems.(5–7) Globally, the prevalence of systemic sclerosis is estimated to be around 10-30 cases per 100,000 population, with increased frequency in the Global North.(8) Despite the significant clinical advances in managing the lethal complications of systemic sclerosis, longer survival with non-lethal complications such as musculoskeletal comorbidities contributes to the overall reduced quality of life in systemic sclerosis patients.(9, 10) Tendon involvement occurs early in the development of systemic sclerosis, and it is one of the strongest predictive factors of overall disease progression.(11, 12) Although the aetiopathogenesis of systemic sclerosis remains poorly understood, a complex picture of genetic susceptibility and self-amplifying dysfunctional repair responses is slowly coming into focus.(9, 13) Nonetheless, relevant insights into molecular mechanisms underpinning tendon involvement are scarce, and further progress is potentially hindered by ethical concerns limiting access to tendon biopsies or scarcity in experimental models that mimic key aspects of the tendon fibrotic niche. (11, 14, 15)

The hallmark of systemic sclerosis is progressive fibrosis that is dominated by persistent activation of resident stromal cells, and excessive deposition and crosslinking of extracellular matrix (ECM). This process is thought to ultimately mediate self-amplifying activation loops accompanied by irreversible replacement of tissue stroma with a rigid, mechanically-strained connective tissue.(16–19) How extracellular matrix mechanical cues contribute to initiation or progression of fibrosis is an area of intense investigation.(19–21) In tendon, repetitive overloading disturbs the underlying homeostatic tension that is critical for maintaining matrix structure and function.(22) Injury responses are intimately linked to cell- and tissue-level mechanical homeostasis, with the extracellular matrix providing biophysical signals for appropriate remodeling and resolution responses.(22–24)

For instance, it is now evident that matrix stiffening precedes the onset of clinical fibrosis.(21, 25, 26) Whereas matrix mechanical cues are instrumental in the successful transition from activated wound healing programs to successful resolution, persistent aberrant matrix stiffening is implicated in instilling fibroblast activation and pathological tissue remodeling.(20) How these processes play out in the context of tendon involvement in systemic sclerosis is unknown.

Engineered biomimetic *in vitro* tissue models provide an attractive avenue to explore the interplay of matrix mechanical cues and fibrosis progression in a precise and controlled manner.(27) The most widely used technique for tuning matrix stiffness while simultaneously characterizing the magnitude of cell-generated forces involve measuring deformations of planar two-dimensional (2D) membranes or 3D hydrogel substrates as a proxy for physical forces.(28, 29) One major limitation of such approaches is the inherent assumption that more deformation translates to more traction forces, in addition to the confounding coupling of several multiple biophysical properties (e.g. stiffness and porosity). An alternative approach is combining synthetic elastic substrates of tunable stiffnesses and traction force microscopy (TFM). In this method, cells are seeded on deformable, non-degradable 2D planar surfaces that are patterned with small trackable markers, *e.g.* fluorescent beads or nanoparticles.(30, 31) Cells and the underlying substrate are sequentially imaged in a stressed (contractile) state, and again after cytoskeletal relaxation. The two images are analyzed to track the beads displacements and to quantify the traction forces that are needed to displace the fluorescent markers. Although the TFM methods are powerful in resolving forces with a subcellular spatial resolution, it requires significant experimental and computational resources that are complex, laborious and often not accessible to laboratories lacking engineering expertise. (32) Furthermore, cells in 2D planar *in vitro* models are highly polarized and often interact with the underlying substrate only on one side while in contact with cell culture medium on the other end.(33)

Hence, such reductionist approaches do not fully capture the complexity between form and function of connective tissues like tendons. In vivo, tendon stromal cells reside in highly aligned 3D niches that are rich in collagens and other proteinaceous components, such as proteoglycans, while being exposed to substantial levels of mechanical tension.

Physically-constrained hydrogels culture models offer an attractive approach to reconstitute this complex mechano-reciprocity of tendon cell-ECM interactions *in vitro*.(34–38) In this approach, stromal cells are embedded in ECM-derived hydrogels that are anchored to rigid pillars in order to geometrically-limit isotropic contractions of the matrix, i.e. uniform compactions in all directions. As a result, cell-generated contractile forces compact the microtissue around the anchoring pillars, deflecting the free ends of each pillar inwards toward the center and driving cell-matrix alignment longitudinally along the axis of principal stresses. The shape of microtissues can be controlled by adjusting the geometry and spacing of the rigid pillars(39), while the tissue contractile forces can be quantified using beam theory or Hooke’s law.(40)

Here, we hypothesized that elevated matrix tension mediates the persistence of fibrotic remodeling in SSc tendons. First, we engineered a modular mechano-culture platform that enables one to easily vary static mechanical tension of tendon-like mimetics, independently of changing bulk tissue properties, thus emulating the complex mechano-reciprocity of tendon structure-function relationship. Next, we characterized tendons involvement in a transgenic mouse model approximating the systemic phenotypes of human SSc and used these models to explore the stromal-immune interactions *in vitro*. We conclude that mechanically-driven, feed-forward loops are likely to be a driving feature of tendon pathology in systemic sclerosis.

## Results

### Development of novel tissue model for tendon mechanobiology

Tendons are mechanically-anchored fibrous tissues, wherein resident stromal cells are aligned longitudinally along the axis of mechanical tension.(41, 42) To recapitulate the tensional state of tendons *in vitro*, we engineered a modular, mechano-culture platform that supports manipulation and monitoring of traction forces generated by 3D tendon-like hydrogel constructs under easily variable mechanical tension (**Figure 1A and B**). We assembled the device by leveraging off-the-shelf components that can easily be sourced by non-engineering laboratories, thus circumventing the need for often inaccessible methods such as complex 3D printing, photolithography or microfabrication techniques.(43) The device consists of two interlocked 12-well plates that are separated by a spacer, and it is accessible from all sides to facilitate media replenishment and gas exchange. The upper plate contains long vertical surgical steel posts, and it locks firmly into an optically clear bottom plate. When the plates are fit together, the two vertical posts in each well are contained within an elliptical hydrogel reservoir. This design feature supports anchoring the 3D tendon-like tissues around the posts, while simultaneously maintaining the tissues in close proximity to the floor of the bottom well for microscopic observations (**Figure 1A**).

**Figure 1.**
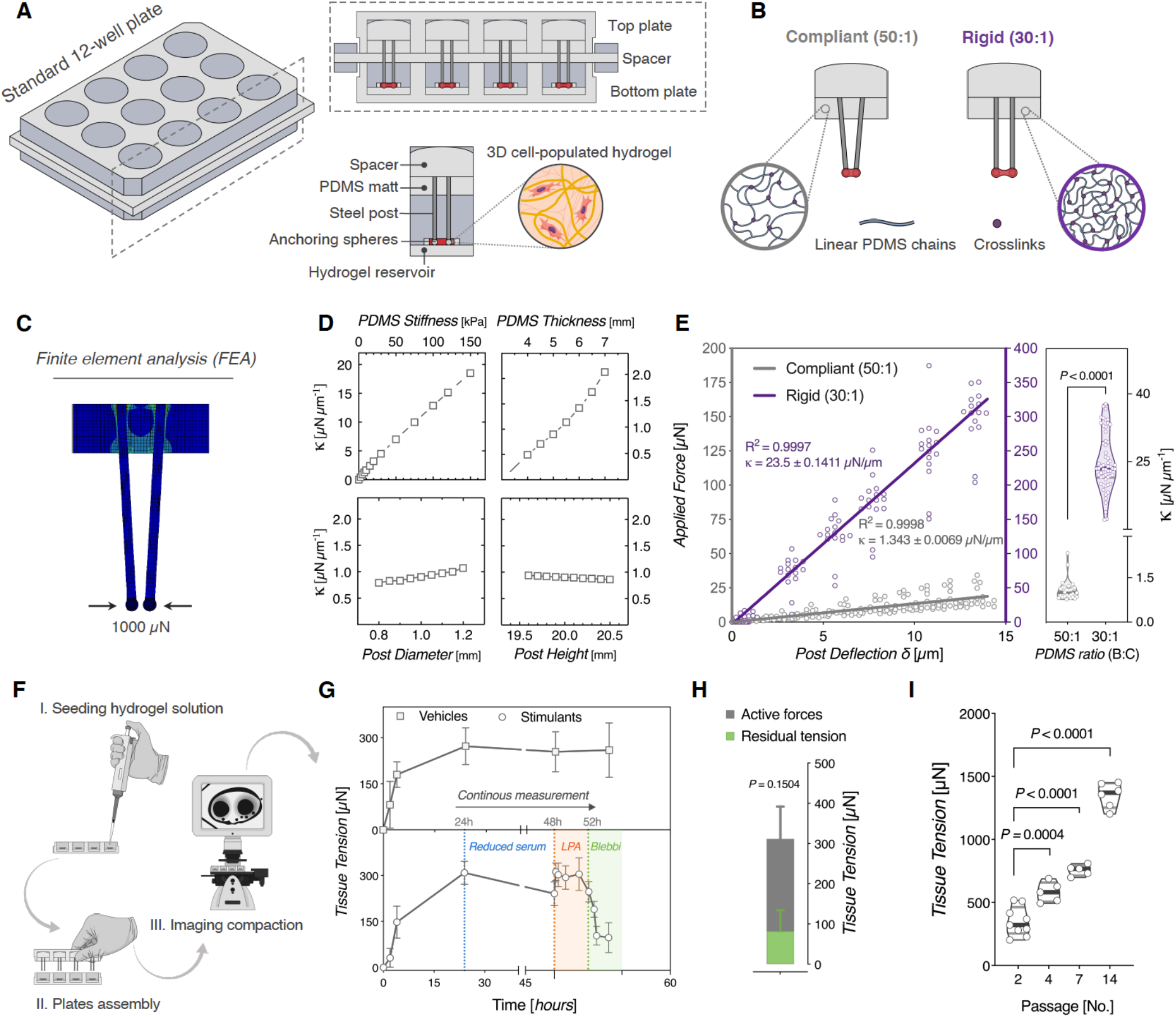
Design and characterization of a modular, mechanically-tunable force sensing platform. **(A)** Schematic illustration of two spaced 12-well plates interlocked on top of each other. Top image: cross-sectional views of steel pillars arrangement in the top plate and hydrogel reservoir within the bottom plate. Bottom image: assembled or precast modular components within each well. **(B)** Mechanical boundary stiffness of 3D tissue-engineered constructs is tuned by varying the mixing ratios of the PDMS siloxane base and curing agent. **(C)** Finite element method modelling of posts deflection in response to applied horizontal force of 1000 µN. **(D)** Post spring constant (*k*) plotted as a function of PDMS stiffness (kPa), PDMS mat thickness (mm), post diameter (mm), and post height (mm), as analyzed by FEM modelling. **(E)** Mechanical characterization of anchoring pillars. Left panel: Force-displacement curves: pillars deflection (*δ*) is plotted as a function of the applied force (*F)*, as measured by a calibrated piezoresistive Femto-Tools™ probe (Solid lines represent linear regression lines of individual data points, *p* <0.0001). Right panel: Violin plots of calculated spring constants (*k*); a measure of steel pillars rigidity (*n* = 58 posts (30:1), 28 posts (50:1), Mann Whitney test, Cohen’s *d* = 4.59 [95.0%CI 3.85, 5.3]). **(F)** Process workflow illustration of tissue formation, plate assembly and imaging of pillars deflection using a standard benchtop microscope. **(G)** Top: Time-course of cell-generated traction forces. Lower: Temporal responses of tissue constructs contractile forces to LPA or Blebbistatin stimulation. **(H)** Active and residual tension at the end of the 48h timepoint of vehicle vs stimulants conditions (*n* = 7-10 tissues, Two-tailed unpaired *t*-test with Welch’s correction, Cohen’s *d* = 0.797 [95.0%CI - 0.278, 1.9]). **(I)** Quantification of tissue tension as a function of the passage number of rat tail-derived tendon stromal cells (*n* = 5-9 tissues, One-way ANOVA with Holm-Sidak *post-hoc* test, Cohen’s *d* = P2-P4 2.24 [95.0%CI 1.13, 3.67], P2-P7 4.23 [95.0%CI 3.0, 6.68]). Estimation plots and permuted *P* values for (E, G, and I) are in (Supplementary figure 2). Cohen’s d effect sizes and CIs are reported above as: Cohen’s d [CI width lower bound; upper bound].

In general, hydrogel stiffness can be modulated by increasing the monomers amounts or by increasing the crosslinking density. However, both approaches eventually manipulate materials bulk properties; i.e. ligand density and presentation, and porosity. To decouple effects of hydrogel rigidity from other bulk properties, we modulated boundary rigidity of tethered collagen hydrogels by varying the base-curing ratios of anchoring PDMS mats (**Figure 1B**). This ultimately allowed for altering effective stiffnesses of hydrogels (i.e. increased resistance felts by cells) while keeping initial collagen amounts and crosslinking density constant. First, we performed numerical simulations to explore which design parameters may have the largest impact on the steel post’s spring constant.

Using finite element (FE) analysis methods, we simulated the posts transverse deflections in response to a 1000 µN force acting on the surface of the beads, while varying post geometries, PDMS mat properties and boundary conditions (**Figure 1C**). The cantilever post was modeled as a stainless-steel material, whereas the bead and PDMS layer were set as linear elastic materials. The FEA model predicted that PDMS stiffness and the active length of the post (i.e. length from the surface of PDMS mat to the post’s free end or in other words the PDMS thickness) have the strongest influence on the deflection of the posts (**Figure 1D**). Since the active length of the post is restricted by the design requirements and well dimensions, we reasoned that varying the PDMS stiffness is the best strategy to modulate the boundary mechanics. Next, we experimentally calibrated the spring constant (*k*) of posts when embedded in compliant (50:1) or rigid (30:1) PDMS mats. Using capacitive force sensors, we derived the spring constants of the posts which is the slope of the force-displacement linear relationship (**Figure 1E** and **Supplementary figure 1**). Rigid posts were approximately 20-fold stiffer with measured spring constant of 23.5 ± 0.14 µN/µm, compared to 1.34 ± 0.0069 µN/µm in compliant posts. These empirically-derived values of spring constants were used to calculate the amount of forces exerted by cell-populated tissues under different mechanical boundaries. To generate tendon-like constructs, we embedded tendon-derived stromal cells in collagen type I hydrogel solution. Cells compacted the polymerized hydrogel into a dense fibrous tissue around the posts anchoring spheres in each well (**Figure 1A and F**). Within hours after seeding, tissue tension evidently increased after two hours, and continued to rise by 3-fold reaching a plateauing maximum of (272 ± 60 µN) at 24 h (**Figure 1G, top**). In that same experiment, we simultaneously tested the diffusion limits of the system by examining rapid dynamics in tissue tension forces in response to stimulation with soluble factors. We found that cell-mediated tissue tension was reduced after the serum starvation, then increased or decreased within approximately 15 min of stimulation with myosin activator, lysophosphatidic acid (LPA), or with myosin ATPase inhibitor, blebbistatin, respectively (**Figure 1G and H**).

Next, we examined the sensitivity of the system in measuring relative differences in tissue tension forces of relatively quiescent or activated stromal cells. We initiated activation of stromal cells by continuously passaging the cells in in plastic tissue culture surfaces which is known to induce long-term mechanical memory.(44, 45)

As we expected, tissue traction force significantly increased over passages by 67% at passage 4 and 117% at passage 7, reaching ∼1353 µN at passage 14 (**Figure 1I**). Taken together, we established a 3D experimental platform comprising of tunable mechanical boundaries that otherwise presented equal bulk material properties to cultured cells.

### Reconstitution of fibrogenesis in tensioned tendon-like constructs

Numerous studies have implicated aberrant TGFβ signaling in the initiation and progression of systemic sclerosis.(46–51) Similarly, degenerative tendon diseases are increasingly viewed as fibroinflammatory pathologies characterized by impaired expression of TGFβ superfamily and dysfunctional scarring.(52–54) TGF-β1 is considered a key profibrotic cytokine that mediates its fibrogenic effects, at least in part, by promoting the activation of resident “quiescent” stromal cells into highly-contractile myofibroblasts.(55–58) Motivated by this, we first sought to examine whether exogenous TGF-β1 stimulation induces activation of naïve tendon stromal cells into a contractile phenotype within tensioned tissue constructs. Untreated stromal cells in reduced serum (i.e. 1% FBS) were used as controls (**Figure 2A**). Continuous treatment with TGF-β1 elicited visible deflections of cantilever posts over a 72-hour period, with a significant two-fold increase in contractile forces reaching a maximum force of 997 ± 141.1 µN at 48 h. (**Figure 2B and C**). Whereas forces continued to rise after 48 h under TGF-β1 treatment, tissue tension plateaued at 24 h at 511 ± 61 µN in the non-stimulated controls. However, these changes were not associated with alterations in metabolic activity, as measured by ATP assay (**Figure 2D**). To correlate functional changes in cellular contractility to phenotypic activation of stromal cells, we performed immunofluorescence staining for α-SMA; a cytoskeletal biomarker indictive of myofibroblast activation. TGF-β1-treated conditions had approximatly three-fold significantly higher fluorescence intensities of α-SMA than those in untreated controls consistent with (**Figure 2E and F**). Furthermore, quantitative analysis of cytoskeletal alignment and nuclear circularity of embedded cells revealed that TGF-β1 stimulation induced higher degrees of cellular alignment and nuclear elongation along the axis of mechanical tension, which reflects an an active remodeling process (**Figure 2G and H**). Collectively, these results underscore to the utility of the mechano-culture platform in recapitulating the hallmarks of pathological fibrosis in tensioned tendon-like tissue constructs.

**Figure 2.**
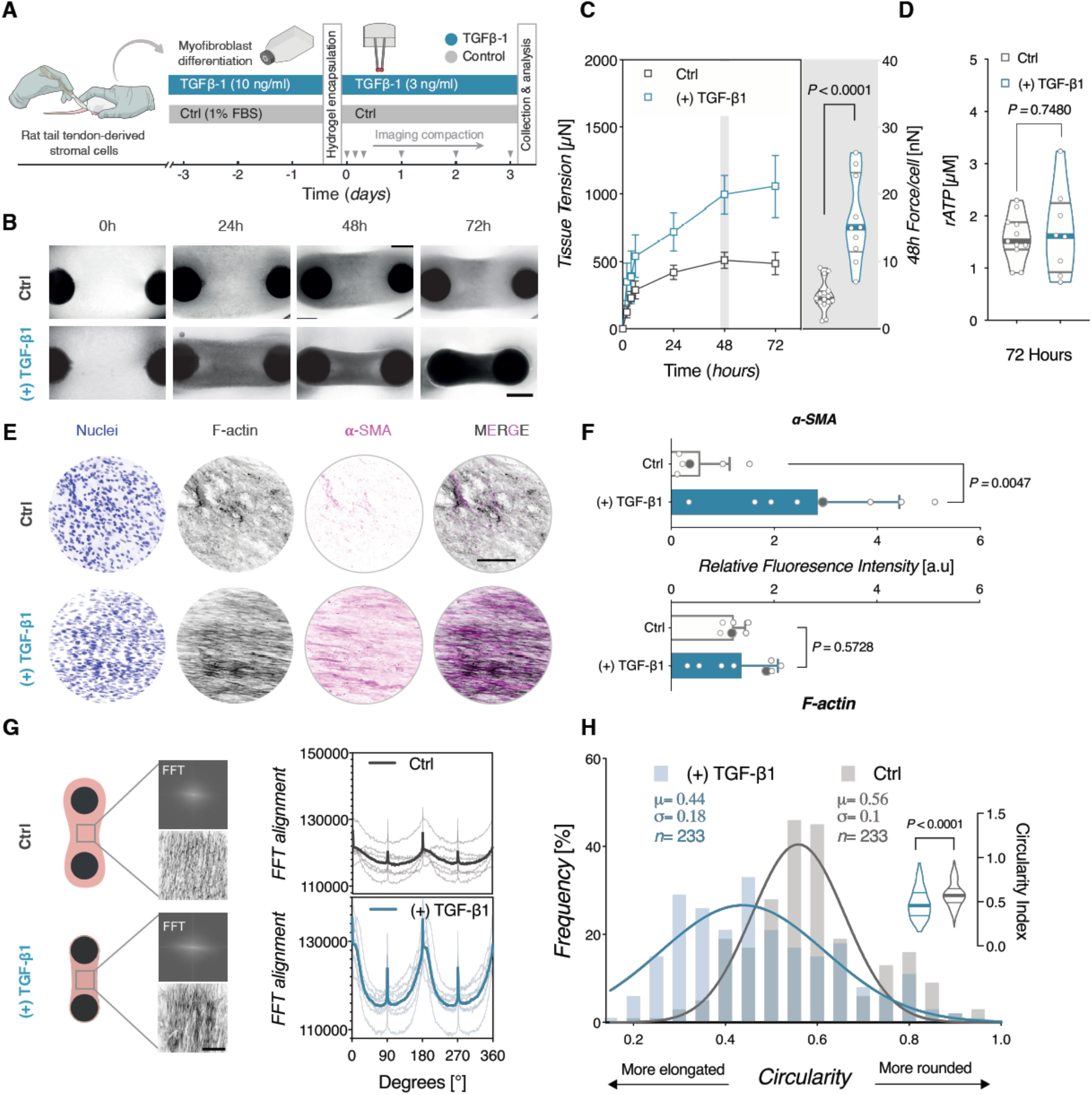
Reconstitution of fibrogenesis in tensioned tendon-like constructs. (**A**) Schematic representation of the experimental protocol. Rat tail tendon-derived stromal cells were pre-treated with 10 ng/ml of TGF-β1 or vehicle control for 72h, before encapsulation in 3D collagen hydrogels. (**B**) Representative images showing the time-course of tissue compaction. Cell-generated forces compact the collagen hydrogel around each post, and continue to displace the posts overtime. Significant posts deflection can be seen in TGF-β1-treated condition. Scale bar = 1 mm. (**C**) Quantitative analysis of tissue traction forces. Left: Evolution of tissue traction forces plotted as a function of time. *n* = 18-21 tissues from 3 biologically independent experiments. Each data point indicates the mean ± SEM. Right: Violin plots of forces per cell, following normalization to initial seeding density. Horizontal lines indicate the median and interquartile range (*n* = 10-15 tissues, Two-tailed Mann Whitney test, Cohen’s *d* = 2.61 [95.0%CI 1.48, 3.52]). **(D)** Analysis of cellular metabolic activity, as a measure of viability. No significant differences were observed between the groups (*n* = 8-11 tissues from 2 biologically independent experiments, Two-tailed unpaired *t*-test with Welch’s correction, Cohen’s *d* = 0.168 [95.0%CI −0.924, 1.21]). **(E)** Representative fluorescence confocal images of cytoskeletal and pro-fibrotic activation markers. Scale bar = 100 µm. **(F)** Quantification of fluorescent intensity levels of F-actin and smooth muscle alpha-actin (α-SMA) in TGF-β1 stimulated and vehicle control. (*n* = 6-8 tissues from 2 biologically independent experiments, Two-tailed Mann Whitney test. α-SMA: Cohen’s *d* = 1.8 [95.0%CI 0.57, 2.93]. F-actin: *d* = 0.276 [95.0%CI −0.822, 1.58]). Grey symbols indicate values for the representative images in **E**. **(G)** Cytoskeletal alignment in TGF-β1 treated and untreated controls. Left: Schematic drawing illustrating approximate positions for image sampling. Insets show representative FFT frequency patterns and corresponding F-actin staining images. Right: Analysis of overall cytoskeletal orientation from FFT graphs indicating the predominant cytoskeletal alignment (Spaghetti plots: *n* = 6 samples (Ctrl), *n* = 7 samples (TGF-β1); solid lines represent the mean values). **(H)** Quantification of nuclear circularity of tendon-derived stromal cells. Frequency distribution analysis shows significant reduction in circularity index in TGF-β1 conditions, indicating more elongated nuclear profiles (*n* = 233 nuclei, solid lines represent Gaussian non-linear fit of frequency values). Inset violin plots represent all individual data points used for the frequency distribution analysis. (Two-tailed Mann Whitney test. Cohen’s *d* = −0.674 [95.0%CI −0.87, −0.48]). Scale bar = 100 µm. Estimation plots and permuted *P* values for (C, D, F, and H) are in (Supplementary figure 2). Cohen’s *d* effect sizes and CIs are reported above as: Cohen’s *d* [CI width lower bound; upper bound].

### Sustained mechanical tension recapitulates key aspects of pro-fibrotic activation in tendon-derived stromal cells

Progressive matrix remodeling in fibrosis results in structural changes (e.g. increased tissue stiffness) that trigger feed-forward, self-sustaining activation loops.(13, 45, 59) We examined how naïve tendon-derived stromal cells would respond to sustained mechanical tension cultured under rigid boundaries, and how mechanical feedback can impact phenotype of tendon-derived fibroblasts (**Figure 3A**). Under rigid boundary stiffness, we observed that cell-generated tissue tension significantly increased by approximately 25-fold after two hours, reaching a maximum value of 6904 µN (SEM ± 1113) after 24 h (**Figure 3B – C**). At 48h, rat tail tendon-derived cells embedded within hydrogels tethered to compliant posts (κ= 1.08 µN/µm) exerted approximately 5 nN per cell [Mean= 5.6 nN ± 2.6, n= 6], while cells anchored to rigid posts (κ= 24.9 µN/µm) had ∼21-fold higher forces [Mean= 117 nN ± 32, n= 7] (**Figure 3C**). Yet, these changes were not associated with significant alteration in cellular metabolic activity after plateauing of tissue tensional forces, despite the profound initial differences in cellular contractility (**Figure 3D**). Next, we investigated whether the sustained mechanical feedback by itself can regulate the phenotype of naïve “quiescent” tendon stromal cells. We first explored the relative impact of boundary stiffness on activation of myofibroblasts. Rigid boundaries induced an approximately 50% increase in α-SMA immune intensities above compliant controls (**Figure 3E and F**), which was similarly consistent at the transcriptional level (**Figure 3G**). Moreover, analyses of cytoskeletal alignment and nuclear orientation revealed higher degree of cellular alignment and nuclear elongation along the longitudinal axis in rigidly tethered constructs (**Figure 3H-I**). Unexpectedly, we found that expression of pro-fibrotic matrix proteins, *Col1a1* and *Col3a1*, is significantly downregulated, whereas *Dcn* is highly expressed under rigid boundaries (**Figure 3J - L**). In similar findings, expression markers indicative of tenogenic linage commitment of stromal cells, (*Scx, Tnmd, Mkx*) were significantly downregulated in highly-tensioned constructs (**Figure 3K**). In sum, these results suggested that sustained mechanical tension can feedback to independently regulate the myofibroblasts pheno-conversion of tendon-derived stromal cells.

**Figure 3.**
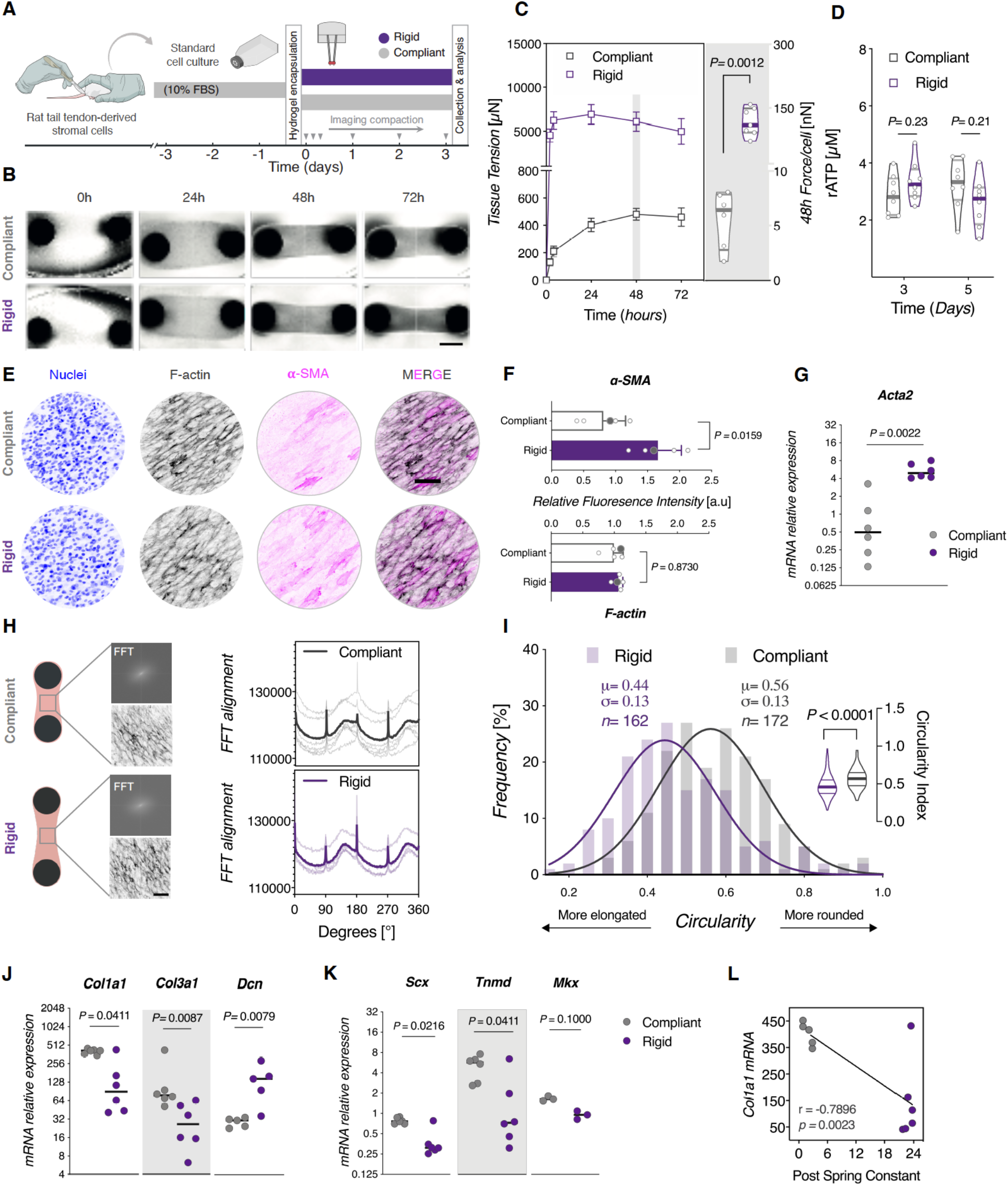
Sustained mechanical tension recapitulates key aspects of pro-fibrotic activation in tendon stromal cells. (**A**) Schematic representation of the experimental protocol. Rat tail tendon-derived stromal cells were encapsulated in 3D collagen hydrogels that were tethered to compliant or rigid posts for up to 3 days. (**B**) Representative images showing the time-course of tissue compaction. Cell-generated forces compact the collagen hydrogel around each post, and continue to displace the posts overtime. Scale bar = 1 mm. (**C**) Quantitative analysis of tissue traction forces. Left: Evolution of tissue traction forces plotted as a function of time. *n* = 14 tissues from 2 biologically independent experiments. Each data point indicates the mean ± SEM. Right: Violin plots of forces per cell, following normalization to initial seeding density. Horizontal lines indicate the median and interquartile range (*n* = 6-7 tissues, Two-tailed Mann Whitney test, Cohen’s *d* = 4.7 [95.0%CI 2.93, 6.31]). **(D)** Analysis of cellular metabolic activity, as a measure of viability. (*n* = 8 tissues/group from 2 biologically independent experiments, Two-way ANOVA (boundary stiffness, time) with Holm-Sidak *post-hoc* test, Cohen’s *d*: Rigid-Compliant (Day 3) = 0.692 [95.0%CI −0.396, 1.7]; Rigid-Compliant (Day 5) = −0.74 [95.0%CI −1.91, 0.413]). **(E)** Representative fluorescence confocal images of cytoskeletal and pro-fibrotic activation markers. **(F)** Quantification of fluorescent intensity levels of F-actin and smooth muscle alpha-actin (α-SMA) as a function of boundary stiffness. (*n* = 5 tissues from 2 biologically independent experiments, Grey symbols indicate values for the representative images in (E). Two-tailed Mann Whitney test. α-SMA: Cohen’s *d* = 2.39 [95.0%CI 0.999, 3.67]; F-actin: *d* = 0.0704 [95.0%CI −0.0358, 0.229]). **(G)** mRNA expression of the pro-fibrotic gene, *Acta2*. (*n* = 6 replicates/group from 3 biologically independent experiments, with each data point representing a ΔCt value of 2-3 pooled tissues, horizontal lines indicate the median, Two-tailed Mann Whitney test). **(H)** Cytoskeletal alignment of tendon-derived stromal cells tethered to compliant and rigid posts. Left: Schematic drawing illustrating approximate positions for image sampling. Insets show representative FFT frequency patterns and corresponding F-actin staining images. Right: Analysis of overall cytoskeletal orientation from FFT graphs indicating the predominant cytoskeletal alignment (Spaghetti plots: *n* = 6 samples (Compliant), *n* = 4 samples (Rigid); solid lines represent the mean values). Scale bar = 100 µm. **(I)** Quantification of nuclear circularity of stromal cells. Frequency distribution analysis shows reduction in circularity index in rigid boundaries, indicating more elongated nuclear profiles (*n* = 172 nuclei (Compliant), *n* = 162 nuclei (Rigid), solid lines represent Gaussian non-linear fit of frequency values). Inset violin plots represent all individual data points used for the frequency distribution analysis. (Two-tailed Mann Whitney test. Cohen’s *d* = −0.671 [95.0%CI −0.903, −0.442]). Scale bar = 100 µm. **(J)** mRNA expression of ECM-related genes, and **(K)** tendon lineage-related genes in stromal cells tethered to different mechanical rigidities. (*n* = 3-6 replicates/group from 3 biologically independent experiments, with each data point representing a ΔCt value of 2-3 pooled tissues, horizontal lines indicate the median, Two-tailed Mann Whitney test). All individual gene expression is shown normalized to *Eif4a2* and *Gapdh* reference genes. **(L)** Scatterplot of expression values for *Col1a1* are plotted against Posts spring constants for linear regression and correlation analysis (Pearson’s r and *p* value are indicated). Estimation plots and permuted *P* values for (C, D, F, I, J and K) are in (Supplementary figures 4 and 5). Cohen’s *d* effect sizes and CIs are reported above as: Cohen’s *d* [CI width lower bound; upper bound].

### Systemic sclerosis-derived tendon cells are autonomously activated independent of matrix mechanics

Fibrotic involvement in systemic sclerosis has been historically investigated in either easily accessible tissues, e.g. skin, or in internal organs that are associated with high mortality rates such as lungs or kidneys. Tendons are often involved early during the disease course, and have high predictive values for further projecting SSc progression over time.(60) Nonetheless, knowledge about how tendons are affected at the molecular level is scarce. This is primarily due to considerable ethical concerns in obtaining research biopsies from a tissue with limited healing capacities and the potential risks of further harming the patients. Here, we give a snapshot of SSc-derived tendon fibroblasts from a rare autopsy that has been donated to our laboratories after a post-mortem examination (only one donor being available for over a period of 4 years).

Next, we assessed the behavior of pathologically activated tendon fibroblasts using our mechano-culture platform to quantify cellular (dys-)functional contractility. For this purpose, we cultured anatomically-matched tendon fibroblasts from the SSc donor or healthy subjects under mechanically compliant boundary conditions (**Figure 4A**). Over a 48-hour period of culture, we observed that SSc-derived fibroblasts showed progressive increase in tissue traction forces similar to normal fibroblasts under TGF-B1 stimulation (**Figure 4B and C, Figure 2C**). At 24h, SSc-derived cells exerted approximately 9 nN per cell [Mean= 8.9 nN ± 1.24, n= 15], while healthy controls had ∼2-fold lower forces [Mean= 5 nN ± 0.76, n= 15]. Next, we focused on characterizing the fibro-inflammatory phenotypes of resident stromal cells populating SSc tendons, and also examined how SSc-derived cells would respond to changes in constructs rigidity. Analysis of mRNA revealed that diseased cells showed a gene expression signature typical of SSc fibro-inflammatory activation, with significant dysregulation of *FN(EDA), IL6, IL8, ACTA2, PDPN and TLR4* mRNA levels (**Figure 4D-F**). Strikingly, we found that *COL1A1* and *COL3A1* mRNA were significantly downregulated in SSc-derived cells compared with anatomically-matched healthy controls (**Figure 4D**). Moreover, *COL1A1* mRNA was significantly downregulated under rigid boundary mechanics to levels comparable to SSc conditions, whereas *ICAM1* and *IL8* expressions were significantly increased in SSc-derived cells in response to dysregulated mechanics. Consistent with our previous observations (**Figure 3L**), *COL1A1* expression in healthy stromal cells showed strong dependency on post rigidity, which was not the case in diseased SSc cells suggesting inability of SSc-derived cells in sensing boundary stiffness (**Figure 4G**). Together, these results demonstrate that SSc-derived tendon cells are intrinsically activated toward hypercontractility in a cell-autonomous manner, with a corresponding absence of matrix stiffness-dependent transcription of ECM collagens and stiffness-sensitivity of stromal-immune interactions markers.

**Figure 4.**
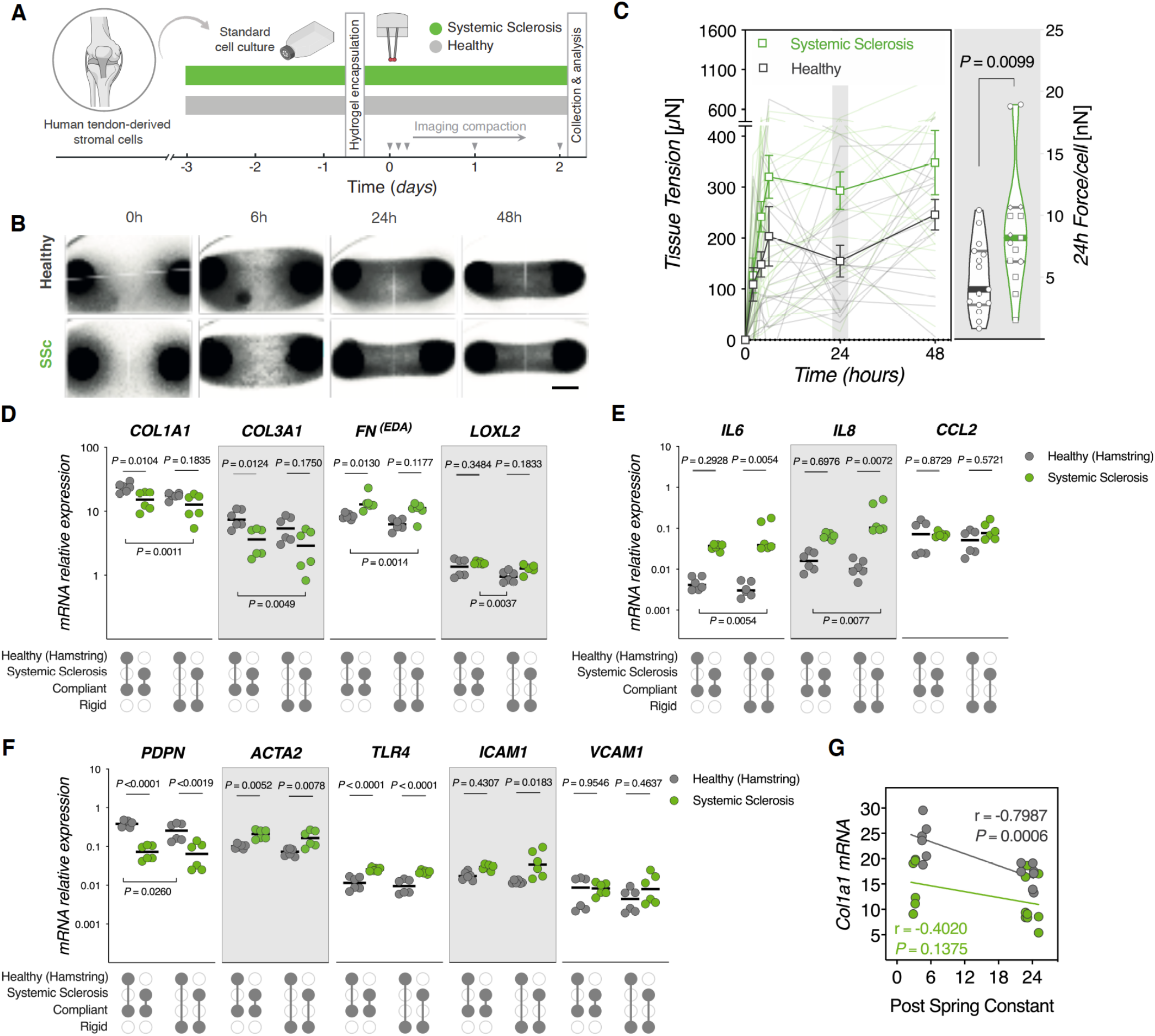
Systemic sclerosis-derived tendon stromal cells are autonomously activated independent of matrix mechanics. **(A)** Schematic representation of the experimental protocol. Human tendon-derived stromal cells from healthy and systemic sclerosis donors were encapsulated in 3D collagen hydrogels tethered to compliant or rigid posts for up to three days. (**B**) Representative images showing the time-course of tissue compaction. Scale bar = 1 mm. (**C**) Quantitative analysis of tissue traction forces. Left: Evolution of tissue traction forces as function of time. *n* = 24 tissues/group from 3 anatomically different tendons of the SSc donor and their matching controls from 3 independent healthy donors. Each data point indicates the mean ± SEM. Right: Violin plots of forces per cell, following normalization to initial seeding density. Horizontal lines indicate the median and interquartile range (*n* = 16 tissues, Two-tailed Mann Whitney test, Cohen’s *d* = 0.928 [95.0%CI 0.305, 1.39]). **(D) (E)** Immunofluorescence and quantification in 2D. **(F)** mRNA expression of ECM-related genes, **(G)** immune/inflammatory genes, and stromal activation markers **(H)** in stromal cells tethered to different mechanical rigidities. (*n* = 6-9 replicates/group from 3 biologically independent experiments, with each data point representing ΔCt, horizontal lines indicate the median, Two-way ANOVA (boundary stiffness, disease stage) with Holm-Sidak *post-hoc* test). All individual gene expression is shown normalized to RPL13A and GAPDH reference genes. **(I)** Linear regression and correlation analysis of *COL1A1* gene expression values and Post’s spring constants (Pearson’s *r* and *p* value are indicated). Estimation plots and permuted *P* values for (C, D, E, and F) are in (Supplementary figures 6, 7 and 8). Cohen’s *d* effect sizes and CIs are reported above as: Cohen’s *d* [CI width lower bound; upper bound].

### Dysregulated mechanics characterize tendon involvement in Fra2^Tg^ mouse model of systemic sclerosis

To extend our findings from the limited clinical samples of SSc tendons, we turned to Fra-2 overexpressing transgenic mouse model which, unlike bleomycin-inducible models, displays spontaneous SSc-like multi-organ inflammatory and autoimmune phenotypes.(61–63) We have recently reported that Fra-2^Tg^ mice develop spontaneous systemic inflammation around 13-17 weeks. The disease course in these mice closely approximates what is seen in human cases with regards to disease onset and progression.(64) Whereas early stages of the disease in mice are dominated by dysregulated T cells activation, late stages are characterized by vasculopathy and extensive multiorgan inflammation and fibrosis (**Figure 5A**).(65)

**Figure 5.**
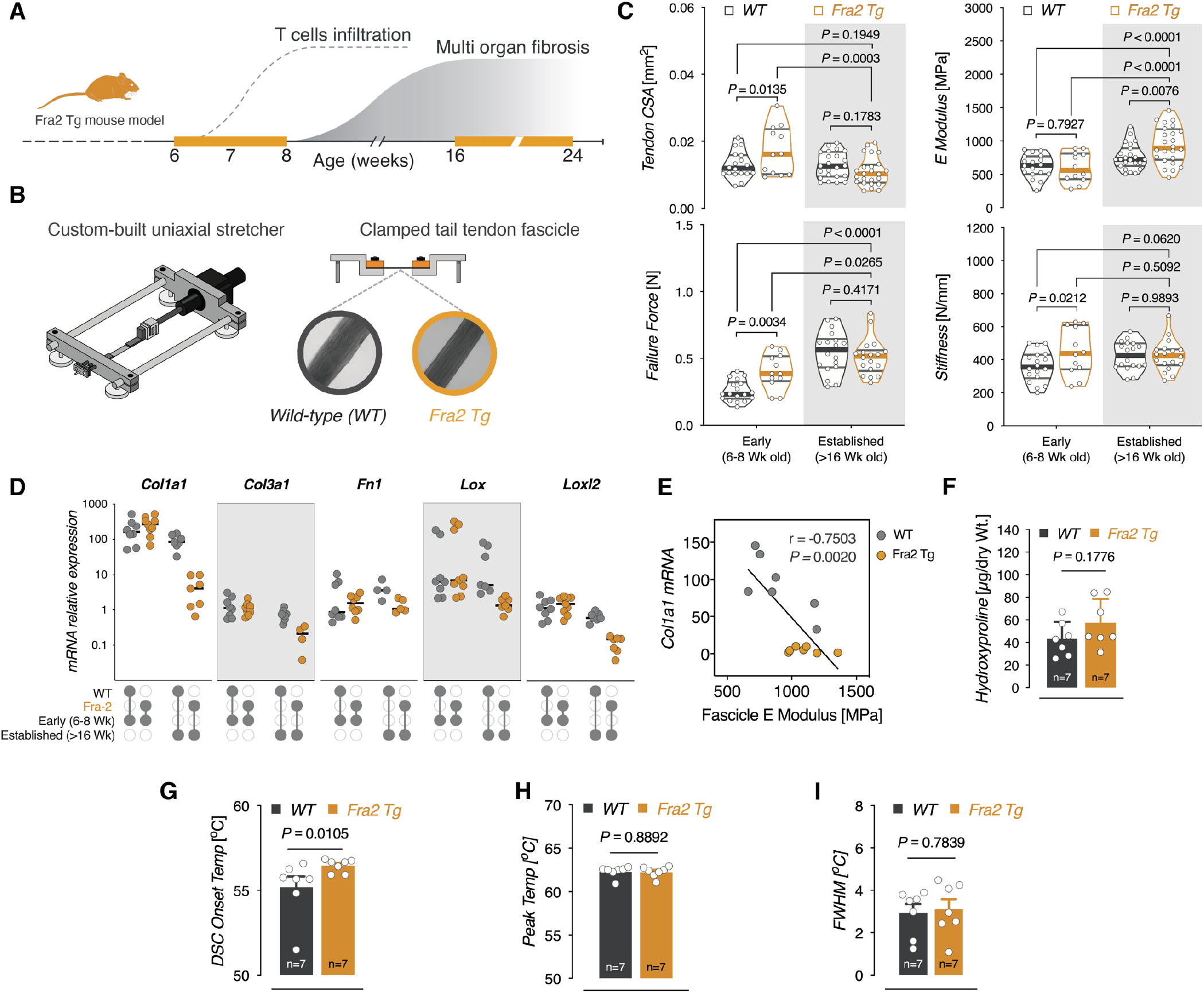
Dysregulated mechanics characterize tendon involvement in Fra2 mouse model of systemic sclerosis. (**A**) Graphical timeline illustrating the natural course of phenotype development in Fra2 transgenic mice. Early-stage disease in Fra2 mice is characterized by multiorgan T cells infiltration (at ∼6-8 weeks), which precedes spontaneous systemic fibrosis at the late-stage of established disease (16-24 weeks). **(B)** Experimental setup for uniaxial mechanical testing. Tail tendon fascicles are clamped in a custom-built stretching device to generate tissue-level stress-strain curves. **(C)** Biomechanical characterization of Fra2 Tg tendons and their WT littermates. Violin plots depict quantification of tail tendon cross-sectional area (CSA), *E* modulus, failure force, and stiffness at early vs. late-stages of established fibrosis. Each data point represents an independent sample; horizontal lines indicate the median and interquartile range (*n* = 12-24 fascicles from 6-7 mice/group, Two-way ANOVA (genotype, disease stage) with Fisher’s LSD *post-hoc* test. **(D)** Baseline mRNA expression of ECM-related genes in fresh frozen Fra2 Tg tail tendons and WT littermates. (*n* = 4-9 mice/group, with each data point representing ΔCt, horizontal lines indicate the median, Two-way ANOVA (genotype, disease stage) with with Fisher’s LSD *post-hoc* test). All individual gene expression is shown normalized to *Anxa5* and *Gapdh* reference genes **(E)** Linear regression and correlation analysis scatterplot of Col1a1 expression in established phenotype are plotted against fascicles *E* moduli (Pearson’s r and *p* value are indicated). **(F)** Quantification of hydroxyproline content in tendons (*n* = 7 mice/genotype, Two-tailed unpaired *t*-test with Welch’s correction, Cohen’s *d* = 0.771 [95.0%CI −0.353, 1.94]). **(G)** Endothermic onset temperature (°C), **(H)** peak temperature, and **(I)** Full-width at half-maximum (FWHM) of thermally-denatured tendons as measured by DSC (*n* = 7 mice/genotype. DSC onset temperature (°C): Two-tailed unpaired *t*-test with Mann Whitney test, Cohen’s *d* = 0.889 [95.0%CI −0.376, 1.48]). Estimation plots and permuted *P* values for (C, F, G, H, and I) are in (Supplementary figure 9 and 10). Cohen’s *d* effect sizes and CIs are reported above as: Cohen’s *d* [CI width lower bound; upper bound].

To investigate tendon involvement in Fra-2^Tg^ mice, we focused our analysis on two cohorts: 6-8 week-old and older than 14 week-old mice; hereafter termed early or established phenotype, respectively. We first characterized the micromechanical properties of tail fascicles, the functional unit of tendons, by performing ramp-to-failure experiments using a custom-made horizontal uniaxial test device (**Figure 5B**).(66) Structural and material properties of tendon fascicles were significantly different in Fra2^Tg^ mice compared to WT littermate controls. While tendon CSA [*P* = 0.0135], failure force [*P* = 0.0034], and stiffness [*P* = 0.0212] were significantly higher during early stages of the disease, mice with established disease displayed appreciably 23% average increase in elastic moduli [*P*= 0.0076] relative to WT littermates **(Figure 5C)**.

Next, we sought to explore potential contributing factors to these differences in tendon mechanics. First, we profiled the expression of a focused set of key matrisome and ECM regulator genes that are known to be differentially expressed in fibrotic pathologies (**Figure 5D**). While relative mRNA expression levels were similar in Fra2^Tg^ and WT in early stage cohorts, expression levels of *Col1a1*, *Col3a1*, *Lox*, and *Loxl2* were significantly reduced in Fra2^Tg^ tendons with established phenotypes. Moreover, *Col1a1* expression showed strong and significant negative correlation with increased *E-moduli* in established phenotypes (*r* = 0.75, *P =* 0.002) (**Figure 5E**).

In contrast, further quantification of total collagen concentration with hydroxyproline assay revealed no differences in average collagen amount (**Figure 5F**). Taking this into consideration, we reasoned that the biomechanical differences in Fra2^Tg^ tendons could emerge from ECM crosslinks, rather than collagen accumulation. To test this hypothesis, we analyzed the molecular thermal stabilities of Fra2 tendons using DSC endotherms. Fra2^Tg^ tendons with established pathology showed significantly higher denaturation temperatures compared to WT controls (T_onset_: *P*= 0.01) (**Figure 5G-I**). This reflects possibly higher degrees of thermal stabilization of collagen molecules. Increased thermal stability suggests that increased collagen crosslinking could underlie the observed mechanical differences, however further analysis would be required to confirm this. Overall, these findings support the premise of disrupted mechano-homeostasis in Fra2 tendons, and highlight a potential transcriptional feedback in the expression of ECM-related genes in response to dysregulated tendon mechanics.

### Dysregulated mechanics and macrophages co-culture differentially regulate stromal fibroblasts activation

In a recent work, we have reported that Fra2^Tg^ mice develop multiorgan systemic inflammation which is associated with increased secretion of pro-fibrotic cytokines IL-10, IL-13, IL-5 and IL-6.(67) Having also observed that rigid boundary stiffness boosted the expression of markers associated with stromal-immune interactions (i.e. IL8 and ICAM1) in SSc-derived stromal cells (**Figure 4**), we sought to investigate whether dysregulated mechanics have an impact on the crosstalk between tendon stromal and innate immune cells. To do so, we used our mechano-culture platform to measure in vitro tissue traction forces of Fra2^Tg^ or WT stromal cells (**Figure 6A**). We cultured tendon-derived stromal fibroblasts from mice with established phenotype under variable mechanical rigidity, either as monocultures or in direct cocultures with bone marrow derived macrophages (BMDM) in a mix-and-match approach (**Figure 6B**). Fra2-derived tendon cells exerted significantly higher traction forces as early as 4hr (*p*= 0.0006), which reached a maximum value of 640.2 ± 104.83 µN at 24hr (*p* <0.0001). The basal tension generated by WT-derived fibroblasts was more than a 4-fold lower than Fra2^Tg^ cells, which leveled off after 6hr in culture (**Figure 6B, left**). In contrast, direct cocultures with BMDM significantly enhanced traction forces generated by WT fibroblasts, with WT BMDM boosting the traction forces by four folds, whereas Fra2 BMDM enhancing WT fibroblasts contractility to levels comparable to activated Fra2^Tg^ fibroblasts (**Figure 6B, Figure 6C**).

**Figure 6.**
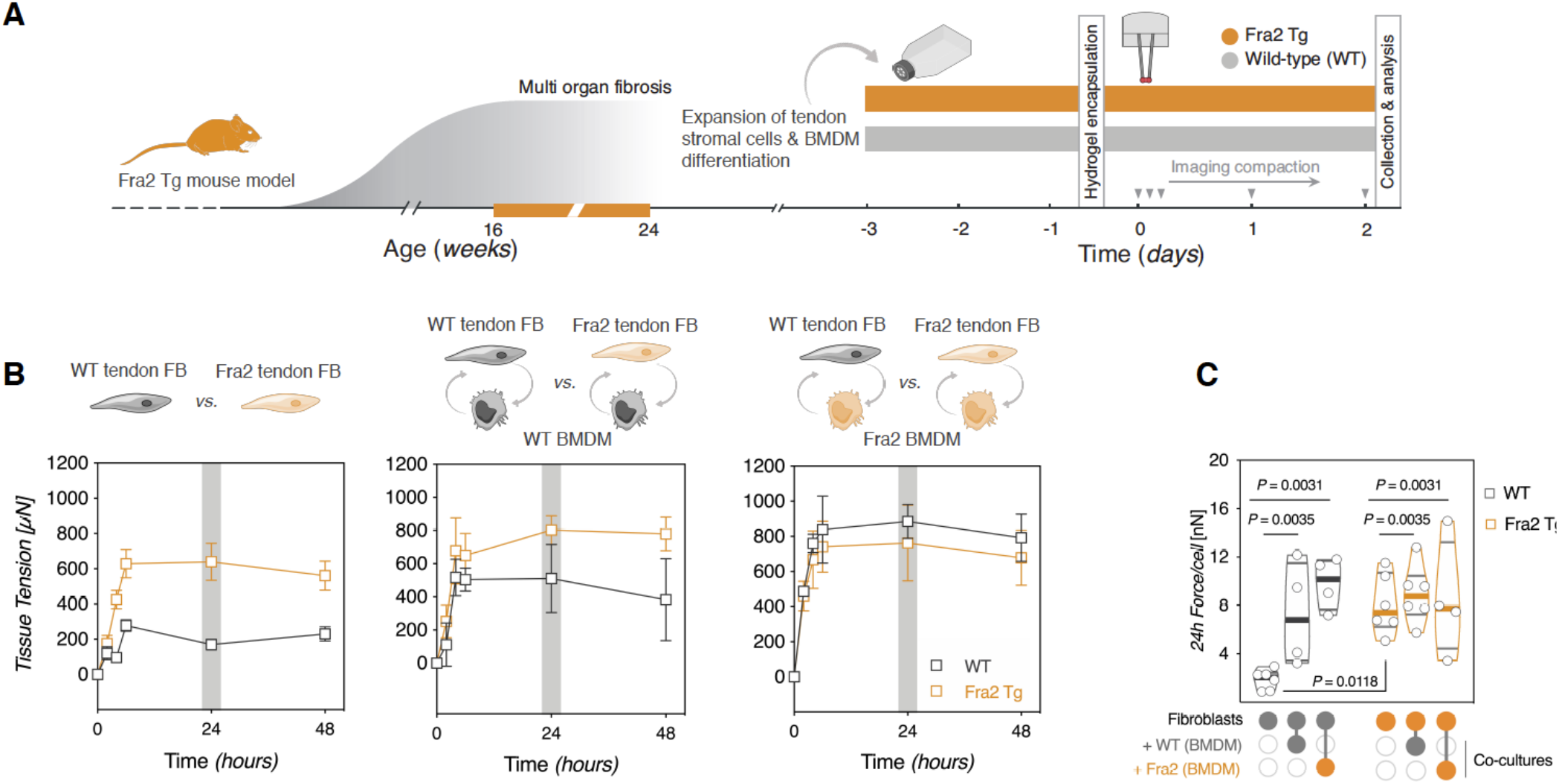
Macrophages enhance stromal activation state of tendon-derived cells. **(A)** Schematic illustration for experimental design and timeline. **(B)** Time-lapsed quantification of tissue traction forces of stromal fibroblasts monocultures (left); stromal fibroblasts co-cultured with WT-derived BMDM (middle); stromal fibroblasts co-cultured with Fra2^Tg^-derived BMDM (right). *n* = 4-6 tissues from 2-4 mice/genotype. Data points represent means ± SEM. **(C)** Violin plots of traction forces per cell, following normalization to initial seeding density of stromal fibroblasts. Horizontal lines indicate median values and interquartile ranges (*n* = 4-6 tissues, Two-way ANOVA (Genotype, co-culture mix) with Tukey’s *post-hoc* test). Estimation plots and permuted *P* values for (C) are in (Supplementary figure 12 A). Cohen’s *d* effect sizes and CIs are reported above as: Cohen’s *d* [CI width lower bound; upper bound].

Next, in a proof-of-concept experiments, we explored how boundary stiffness and cocultures with BMDM influence the expression of profibrotic matrisome and ECM regulator genes in Fra2^Tg^ or WT cells. In monocultures, Fra2^Tg^ tendon stromal cells showed up to two-fold increase in *Col1a1* and *Lox* expression relative to WT controls (**Supplementary figures 11 A-C**). However, this effect was significantly reduced (*Col1a1 P*= 0.042, *Lox p=*0.01) under rigid boundaries mirroring our observations in Fra2 tendons *in vivo* and SSc-derived tendon fibroblasts (**Figure 5D-E and Figure 4F-I,** respectively). Interestingly, we observed consistent trends in the increased relative expression of *Col1a1*, *Col3a1* and *Lox* in co-cultures with differentiated macrophages, irrespective of boundary mechanics (**Supplementary figure 11 D-F**).

## Discussion

We have engineered a new modular mechano-culture platform that enables one to easily vary static mechanical tension of tendon-like hydrogel constructs independently of bulk ECM stiffness, crosslinking density, or composition. We have used this platform to reveal a complex interplay between sustained mechanical tension, ECM remodeling and activation of tendon-derived stromal cells in models of fibroinflammatory pathologies. Fibrotic progression in systemic sclerosis is accompanied by substantial changes in ECM mechanics, particularly heightened matrix tension and rigidity. Until now, a myriad of 3D *in vitro* tissue mimetics has been developed to model states of static tension in connective tissues. However, the artisanal craftsmanship required to engineer and manufacture these systems has limited widespread adoption in biomedical and translational research labs. Our mechano-culture platform described here addresses this need. We propose a straightforward platform that can be easily replicated, allowing reproducible study of pathological fibrosis, the role that matrix mechanics plays in this process, and simultaneously providing a highly relevant online (non-invasive) readout of cellular forces *in situ*.

In the present study, we show that elevated matrix rigidity in 3D tissue constructs mediates phenoconversion of naïve tendon-derived stromal fibroblasts towards activated myofibroblasts. By anchoring collagen tissues to cantilevers posts with varying degrees of boundary rigidity, we sought to create ECM mimetics with states of high or low tension, independently of bulk stiffness of the provided collagen matrix or exogenous application of TGF-β1. In general, tissue resident stromal cells are able to sense different degrees of matrix tension through a sophisticated mechanotransduction apparatus that connects the extracellular matrix to the contractile cytoskeleton and other mechanically sensitive subcellular structures. In high tension environments, this process leads to incorporation of ⍺-SMA into actin stress fibers; a hallmark of myofibroblasts activation.(68) However, most of these observations are based on experiments conducted on 2D planar substrates where mechanotransduction dynamics are substantially different from 3D counterparts. Nonetheless, our findings are consistent with the role of matrix tension in myofibroblasts activation. (68, 69)

How elevated matrix tension contributes to myofibroblasts activation and fibrosis progression is an area of intense research. Elegant work from Wipff *et. al.* and Wu and colleagues have clearly demonstrated that excessive mechanical tension sets resident stromal cells on a fibrotic trajectory by mediating the activation of latent TGF-β signaling loops.(70, 71) Because myofibroblastic activation is associated with a phenotypic switch and enhanced production of extracellular matrix, we focused our analysis on the transcription of key ECM molecules and markers of tendon lineage specification. Strikingly, dysregulated matrix mechanics, i.e. increased tissue stiffness, was associated with downregulation of collagens mRNA transcription in tensioned 3D tissue cultures *in vitro.* This finding is in contrast with previous work using 2D planar substrates to model stiffness-mediated fibrogenesis, in which progressive stiffening strongly promoted fibroblasts matrix synthesis.(72) Using 2D polyacrylamide substrates, we previously showed that increased substrate stiffness elicited transcriptional response in bone marrow mesenchymal stromal cells similar to our findings here with strong downregulation of markers of matrix synthesis and tenogenic differentiation.(73)

In this study, we extended the utility of this mechano-culture platform to investigate the impact of the mechanically stressed niches on the pathological activation of tendon fibroblasts in systemic sclerosis. We provide pilot evidence in proof-of-principle experiments using multiple tendons from a single donor with confirmed diagnosis of systemic sclerosis. Although we acknowledge the limited generalizability of these findings due to the lack of biological replication, to our best knowledge this is the first demonstration of tendon involvement in systemic sclerosis at the cellular and molecular levels. We found that tendon stromal cells from healthy donors exhibited comparable response to rodent-derived rigidly-anchored quiescent fibroblasts *in vitro* and stiff Fra-2^Tg^ tendons *in vivo*. Specifically, healthy tendon-derived cells showed strong downregulation of ECM collagens (*COL1A1* and *COL3A1*) in response to mechanically tensioned rigid posts. In contrast, tendon cells from the SSc patient showed a disordered response to discriminate niche stiffness. Our findings are in agreement with recently published reports showing that dense, mature fibrotic niches were associated with decreased transcription of *Col1a1* and *Col1a2*.(74)

This was further confirmed in a multi-omics single cell profiling of the aging lung which attributed the dysregulated matrix remodeling in aged lungs to the increase in transcriptional noise and aberrant epigenetic control.(75) Although the beforementioned studies attributed these age-dependent effects (at least in part) to cell-mediated processes, the contribution of ECM stiffness, which is known to increase with aging, cannot be ruled out.(76) Based on our findings, it is tempting to speculate that increased matrix stiffness is a potential key driver of dysregulated matrix remodeling in tendons, which was recently shown to be the case in brain tissues.(77)

In strong support of the valuable but limited human data we present, we found that tendons of Fra2 mice exhibit dysregulated mechanical homeostasis, with significantly higher failure forces and elastic moduli. We have recently showed that Fra2 overexpressing transgenic mice develop T cell-mediated spontaneous systemic inflammation with many features reminiscent of systemic sclerosis. Moreover, sera for Fra2^Tg^ mice had elevated levels of profibrotic Th2 cytokines (IL-5, IL-6 and IL-10) as well as other inflammatory cytokine/chemokine mediators (IL-1b, TNF-a and CCL2). Biochemical and structural analyses revealed that tendon ECM stiffening in Fra2 Tg mice likely results from matrix crosslinking rather than excessive ECM production. These findings both mirror and contrast with previous reports examining the role of Fra-2 in tissue fibrosis. In agreement with our findings, Ucero *et. al.* showed that Fra-2^Tg^ mice develop spontaneous lung fibrosis which was associated with increased expression of matrix crosslinking enzyme *Loxl2*.(63) Although the spontaneous fibrosis in Fra-2 lungs was also associated with enhanced expression of ECM collagen, this was not the case in our tendons analysis suggesting tissue-specific differences in the mechanism of fibrosis. Similarly, Georges and colleagues demonstrated that tissue stiffening preceded collagen deposition in an inducible model of liver fibrosis through a Lox-mediated crosslinking activity that ultimately results in myofibroblast activation.(25) Whether tendon stiffening in Fra-2^Tg^ mice is mediated through intrinsic cellular activation, profibrotic Th2 paracrine signaling or both remain elusive and is ground for future work.

However, culturing Fra-2 tendon-derived stromal cells under tension revealed that these cells are highly contractile and express significantly higher levels of *Col1a1* and *Lox* under compliant rigidities. This suggests that tendon stiffening in Fra-2 mice is mediated, at least in part, by resident fibroblasts in a cell-autonomous manner.

We conclude that tension-regulated positive feedback loops are likely to be a central feature of tendon fibrosis. Our study demonstrates how mechanovariant tissue engineered models provide a powerful platform for unwinding the mechanisms of cell-matrix cross-talk. We suggest that these experimental models can and should play an important role in future research, including the identification of novel therapeutic targets to potentially mitigate the onset and progression of fibrosis.

## Methods

### Device assembly

The device was assembled by combining two standard 12-multiwell plates interlocked on top of each other. Plates were separated by a Poly(methyl methacrylate) spacer. The top plate contained two arrays of steel pins pairs (A20007120, inox-schrauben.de) with a diameter of and length of embedded within a thick mat of PDMS. Each well in the top plate’s; the upper (A1-A4 wells) and lower (C1-C4 wells) arrays contains one pair of pins (i.e. 8 pairs in total per plate), while the central wells were left empty. The PDMS layer was casted on top of a of polycarbonate disc. This allows for controlling the PDMS thickness as well as the depth of pins into the bottom plates. Pins were (1mm ± 0.006mm) mm in diameter, (20 ± 0.5 mm) mm in length and spaced approximately 4mm apart from each other mm. The corresponding bottom plate encompasses a PDMS well which serve as reservoirs for casting cell-laden hydrogels. The 5mm x 8mm PDMS reservoirs wells were punched out of ∼2mm thick layer using a metal puncher and glued to a ∼10mm mm of 10:1 PDMS.

### Calibration of steel pillars spring constants

Posts spring constants were measured experimentally using a capacitive piezoresistive MEMS force sensor mounted on an XYZ micro-manipulating arm in a horizontal configuration (FT-RS1002 Microrobotic System: FemtoTools AG, Switzerland), as described elsewhere.(78, 79) The posts were individually calibrated by bringing the tip of the force sensor into contact with approximately the center of the glued spherical beads. Displacement force is measured while simultaneously pushing against the sphere to deflect the post. For each measurement, the post was displaced until the sensor force reached a value of 400µN or 4000µN for the compliant or rigid configurations, respectively. Hooke’s law is used to calculate the spring constant (κ):

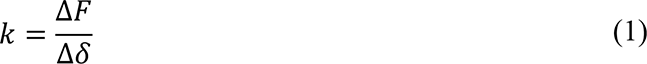

### Quantification of traction forces

To quantify tissue traction forces, cell-populated gels were cultured under tension for the indicated time periods outlined in figures legends. Timepoint “zero” was defined as the moment the growth media is added to the polymerized collagen gels. Briefly, tissue tension was quantified by monitoring the post deflection every 2h for the first 4-6 hours and then every 24h for up to 3 days. Images of the beads capping the posts were acquired with an inverted, brightfield EVOS™ XL Core Imaging System (AMEX1000, Thermo Fisher Scientific) using a 2x or 4x objectives. These images were processed with a custom-written Matlab script (Matlab^®^ R2018b, MathWorks, Inc.) to segment the two beads and calculate the distance between the centroids (in pixels). The deflection of the posts was determined for each time point and the distance between the two beads was rescaled to the initial inter-post distance (in µm) that was experimentally measured with the FemtoTools during the calibration spring constants. Finally, tissue traction forces were calculated with Hooke’s law formula using the measured deflections (Δ*δ*) and the average of spring constant of each post (κ).

### Finite element simulation of microtissue contractility

Posts and the embedding PDMS mat model was implemented in Ansys Workbench (V. 16.2., ANSYS, Inc). The posts were modelled as stainless-steel material using the default settings of the software. spherical beads were simulated as a linear elastic material with a density of 2400 kg/m3, a Young’s modulus of 185 GPa, and the PDMS was modeled as a linear elastic material with a fixed Poisson’s ratio of 0.499 and a Young’s modulus of 5 kPa. A hexahedral mesh was used for the post and PDMS substrate while tetrahedral mesh was fitted for the capping bead. To save computational power, only one post was modelled, and a symmetry region was added create the second post. For the boundary conditions, fixed constrains on all sides were chosen. As loading condition, a force acting on the surface of the beads in direction of the plane of symmetry was set. The magnitude of the force acting as loading conditions was set to 1000 µN. This force represents the force generated by the contracting cells in the collagen gel. The simulations were also done using a Neo-Hookean model for the material.

### Fosl2/Fra-2^Tg^ mice experiments

Fosl2^Tg^ mice were generated by Sanofi-Genzyme as described elsewhere.(67) Briefly, a vector containing the murine Fosl2 gene (Exons 1 to 4, corresponding introns, and truncated UTRs) was randomly inserted into the genome under the control of the MHCI promotor H2Kb. The eGFP sequence was inserted in frame after the Fosl2 transgene as a marker with a self-cleaving peptide T2A (Thosea asigna virus 2A) inserted between the two coding sequences. Experiments with the Fra-2^Tg^ mice were approved by Cantonal Veterinary Office Zurich.(67) Mice were housed under specific pathogen-free conditions at the University of Zurich.

### Biomechanical testing of tendons

For biomechanical testing of tendons, mice were anesthetized and sacrificed according to the guidelines of the Swiss Animal Welfare Ordinance (TSchV) University of Zurich. Samples were then stored in a freezer until further analysis. On the day of mechanical testing, fascicles were extracted by holding the tail from its posterior end with surgical clamps, and gently pulling off the skin until bundles of fascicles were exposed. Each fascicle was examined under a microscope (Motic AE2000, 20x magnification) for visible signs of damage, and for measuring the external diameter. Specimens with frayed ends, visible kinks or diameters less than 90µm were discarded. Micromechanical tensile testing was performed using a custom-made horizontal uniaxial test device to generate load-displacement and load-to-failure curves (10N load cell, Lorenz Messtechnik GmbH, Germany). Briefly, two-centimeter fascicle specimens were carefully mounted and kept hydrated in PBS, as previously described (Snedeker et al., 2005). Each fascicle underwent the following protocol: pre-loading to 0.015 N (L0: 0% strain), 5 cycles of pre-conditioning to 1% L0, an additional 1% strain cycle to calculate the tangential elastic modulus, and then ramped to failure to 20% strain under a predetermined displacement rate of 1 mm/s. The load-displacement data were processed using a custom-written Matlab script (Matlab® R2018b, v. 9.5.0.944444, MathWorks, Inc.). Tangent elastic moduli were calculated from the linear region of stress-strain curves (0.5-1%). Nominal stress was estimated based on the initial cross-sectional area. Cross-sectional area was calculated from microscopic images assuming fascicles have perfect cylindrical shape.

### Isolation of tendon-derived stromal cells from healthy and systemic sclerosis tendons

Hamstring tendons (Semitendinosus and Gracilis) were collected from otherwise healthy donors undergoing surgical autograft repair of their anterior cruciate ligaments.(80) Samples were collected with informed donor consent in compliance with the requirements of the Declaration of Helsinki, the Swiss Federal Human Research Act (HRA), and Zurich Cantonal Ethics Commission (Approval number: 2015-0089). Tendons were immediately placed in (DMEM)/F12 medium (D8437, Sigma-Aldrich) in the operating room and subsequently processed in the cell culture lab. Human tendon-derived stroma cells were isolated by collagenase digestion as described elsewhere.(81) Briefly, tendon samples were washed in PBS to remove blood or tissue debris. Surrounding fat, fascia or muscles were excised, and tendon tissue was cut into approximately 5mm x 5mm pieces followed by 6-12h digestion with Collagenase, Type I (17018029, Gibco™) at 37 °C. Isolated cells were allowed to grow in DMEM/F12 supplemented with 20% heat-inactivated fetal bovine serum (FBS - 10500, Gibco™), 1% (v/v) Penicillin-Streptomycin (P/S, P0781, Sigma-Aldrich) and 1% (v/v) Amphotericin B (15290018, Gibco™). Cells were maintained in a culture incubator set at 37 °C and 5% CO2, and fresh media were replenished every 3-4 days until cells reached 80% confluency in T75 culture flasks. Cells were routinely sub-cultured once during initial expansion, and were used between passages 2 and 5 for all experiments.

### Isolation of tail tendon-derived stromal cells

Tail tendon-derived stromal fibroblasts were isolated from sexually mature 12 to 14 week-old Wistar rats (Zurich Cantonal Veterinary Office permissions: ZH265/14 - ZH239/17), or Fra2 transgenic mice and their wild-type controls. Rat tail tenocytes were chosen as a model in proof-of-concept experiments because they originate from the same population of tendon somatic progenitor cells as load-bearing tendons (82), and are convenient in obtaining a large number of cells at low passages. After euthanasia, tail tendon fascicles were extracted under sterile conditions as described above in the biomechanical testing section. (83) Tissue was washed once in PBS, minced into small pieces, and digested overnight in 0.2% (wt/v) Collagenase D (11088866001, Roche) in DMEM-F12 medium supplemented with 1% (v/v) Penicillin-Streptomycin.

### Isolation and differentiation of bone marrow-derived macrophages

Bone marrow-derived macrophages were prepared as detailed elsewhere.(84) In brief, bone marrow cells were harvested by flushing femurs and tibias of sex-matched, littermate mice. Bone-marrow-derived macrophages (BMDMs) were generated in DMEM - high glucose media supplemented with 10% (v/v) heat-inactivated FBS, 1% (v/v) Penicillin-Streptomycin, and 50 ng/ml of recombinant murine M-CSF (315-02, PeproTech). Half of culture medium was replenished every 3-4 days. At day 7 postinduction, macrophages were collected and used in the co-culture experiments with tendon-derived stromal fibroblasts.

### Cell culture

For all experiments, cells were maintained in Dulbecco’s Modified Eagle’s Medium with L-glutamine, sodium pyruvate, and sodium bicarbonate (DMEM high glucose - D6429, Sigma-Aldrich), supplemented with 10% FBS, 1% Penicillin-Streptomycin, 1% MEM Non-essential Amino Acid Solution (M7145, SAFC Sigma-Aldrich) and 200 μM L-ascorbic acid phosphate magnesium salt (013-19641, FUJIFILM Wako Chemicals).

### Fabrication of cell-laden, tendon-like constructs

Tendon-derived cells were encapsulated in hydrogels by mixing with neutralized collagen solutions prior to gelation, and cultured for 3-5 days. The two time-points were chosen to allow for full compaction of constructs (day 3) and additional days for tissue maturation (day 5). Collagen gels were produced using rat tail collagen type I (High Concentration type I collagen - 354249, Corning^®^). A 2x acidic collagen solution was neutralized on ice to pH 7.2-7.4 with precooled 10× PBS (70011044, Gibco^™^) and 1M sodium hydroxide (S5881, Sigma-Aldrich). Next, this solution was mixed at ratio of 1:1 with a 2x concentrated cell solution to reach a final collagen concentration of 1.7 mg/ml. A 90 µl of the liquified cell-laden gel solution was added per PDMS reservoirs and was allowed to polymerize for 45 min, after assembling the plates. Plates were then carefully removed from the incubator and 1 mL of supplemented culture medium was added to each well. When indicated, the cells were treated with recombinant TGF-β1 (580702, BioLegend^®^), as specified in figure legends, and were cultured in growth media containing 1% FBS.

### RNA extraction from human and Fra2^Tg^ mice tendons

Tail fascicles were placed in 2mL Eppendorf Safe-Lock tubes, snap frozen in liquid nitrogen and stored at −80C until further assayed. On the day of isolation, tubes were placed immediately in dry ice. Samples were transferred to supercooled Spex microvial cylinder (6757C3, SPEX™ SamplePrep) containing 150µl of GENEzol™ reagent (GZR200, Geneaid). Samples were cryogrinded in a bath of liquid nitrogen using FreezerMill (6870, SPEX™ SamplePrep) until fascicles were completely disrupted. Pulverized samples were collected by rinsing the tubes with 900µl GENEzol™, transferred to a clean 1.5 ml tube and kept in dry ice. To purify RNA, tissue homogenate was thawed and mixed well with 200µl of Chloroform (102445, Merck) at a mixing ratio of 1:5. Samples were then spun down for 15min (at 15,000g, 4°C). The upper RNA-containing aqueous phase was transferred to a clean 1.5 ml tube and mixed with one part of 70% ethanol. RNA cleanup was subsequently performed using PureLink™ RNA Micro Scale Kit (12183016, Invitrogen™), including an on-column DNA digestion step with DNase I (DNASE70, Sigma-Aldrich). For collagen gels, constructs were collected in Qiagen RLT Plus lysis buffer (1053393, Qiagen), and were snap-frozen in liquid nitrogen and stored at −80°C. Samples were disrupted by vertexing for 1 min. followed by homogenization using QIAshredder columns (79654, Qiagen). Total RNA was isolated using RNeasy Plus Micro Kit (74034, Qiagen), including a step to remove genomic DNA (Qiagen gDNA Eliminator spin columns). Cleaned RNA concentration and quality were determined by spectrophotometric determination of A_260_/A_280_ ratio. Samples were stored at −80 °C, and only samples with A_260_/A_280_ ratio of 1.8-2 were used for downstream RT-qPCR analysis.

### Real-time quantitative PCR (RT-qPCR)

Real-time quantitative PCR were performed using StepOnePlus Real-Time PCR System (4376600, Applied Biosystems^™^) and TaqMan Assays. 90ng of total RNA were reverse transcribed to cDNA using High-Capacity RNA-to-cDNA Kits (4387406 or 4368814, Applied Biosystems™). RT-qPCR reactions were run with 2μl cDNA and 8μl of TaqMan® Mastermix (containing 5 μl Universal PCR Master Mix, 0.5 μl of TaqMan primer, 2.5 μl of ultrapure water) adding up to a total volume of 10μl.

Primer details and assay IDs are listed in the online supplementary material (**Supplementary Table S1**). Reactions were carried out in technical duplicates. Relative expression was calculated using the comparative 2^-ΔCT^ method.(85)

### Pharmacological treatments of tissues with agonists and inhibitors

To test the dynamic contractility and temporal response to stimulation with soluble factors, cell-laden collagen hydrogels were formed and cultured in standard growth media for 24h as described above. The then constructs were serum-starved in 1% FBS for another 24h to reduce the basal levels of cytoskeletal tension. Cellular contractility was modulated by incubating tissues with Oleoyl-L-α-lysophosphatidic acid (LPA) at a final concentration of 20µM (L7260, Sigma-Aldrich), or (−)-Blebbistatin at a final concentration of 50µM (B0560, Sigma-Aldrich). Vehicle-alone controls were: PBS for LPA and DMSO for Blebbistatin in serum-reduced media.

### Immunostaining and confocal fluorescence microscopy

Collagen constructs were fixed while still under tension with pre-warmed 4% neutral buffered formaldehyde solution (ROTI^®^Histofix 3105.2, Carl Roth) for 45 min at room temperature. Subsequently, tissues were permeabilized with 0.5% Triton X-100 (93418, Sigma) in PBS for 30 min, and blocked overnight with 3% bovine serum albumin (BSA - P6154, Biowest) in a humidified chamber at 4 °C. Samples were immersed in Image-iT FX Signal Enhancer blocking solution (I36933, Invitrogen™) for at least 30 min before overnight incubation with primary antibody solution at 4 °C. After washing, tissues were incubated with the appropriate fluorescently-conjugated, isotype-specific secondary antibody for 4 hours, and protected from light. Nuclei and actin filaments were counterstained with NucBlue and AlexaFluor-conjugated phalloidin, respectively. Samples were mounted in 120 µm Secure-Seal™ adhesive spacers and embedded in Agarose, low melting point (V3841, Promega). Immunofluorescence images were acquired with the iMic spinning disk confocal microscope, fitted with a Hamamatsu Flash 4.0 sCMOS camera and a SOLE-6 Quad laser (Omicron), using 10x (N.A. 0.4) and 20× (N.A. 0.75) objectives (Olympus UPLSAPO).

## Statistics

All statistical analyses were performed using Prism 9.0 (v. Version 9.1.0, GraphPad Software) and DABEST-Matlab for the estimation statistics (Matlab® R2018b, MathWorks, Inc.). Samples were checked for normality distribution with Shapiro–Wilk test. Statistical differences were evaluated with Unpaired Student t-test with Welch’s correction or ordinary one-way ANOVA (multiple comparison), for normally distributed data. Mann–Whitney test was used in cases where the normal distribution criterion was not met. Exact *P*-values and *post-hoc* tests are reported in the figure legends. Whenever applicable, we also report the magnitude of the effect (Cohen’s *d* effect size) and CIs. All the figures for estimation statistics and permutation p-values are in the Supplementary Information.

## Acknowledgments

We are grateful for the donors who generously consented to use their tissue in this work. We are grateful for the support of clinical teams, research nurses, administrative (Ms. Helen Strebel) and the cleaning staff at Balgrist University Hospital and the Swiss Center for Musculoskeletal Biobanking. We would like to thank Mr. Silvio Broder for his assistance with the biomechanical testing, and to Dr. Astrid Jüngel and Dr. Stefan Dudli for the constructive discussion and feedback. This work has been funded by the Cariplo Foundation [2016-0481], the Vontobel Foundation, and institutional funding of both the ETH Zurich and the University Hospital Balgrist.

## Author contributions

A.A.H and J.G.S conceived the study and designed the experiments. R.K, J.F, and A.A.H. prototyped and tested the mechano-culture platform. A.A.H analyzed and interpreted the data and wrote the manuscript. F.R. assisted with Fra2 mouse experiments. S.L.W. performed biomechanical tests. B.N assisted with experimental work and performed gene expression experiments. O.D provided resources. J.G.S acquired funding, supervised the project, interpreted the data, and commented on the manuscript.

## Supplementary Information

**Supplementary figure 1.**
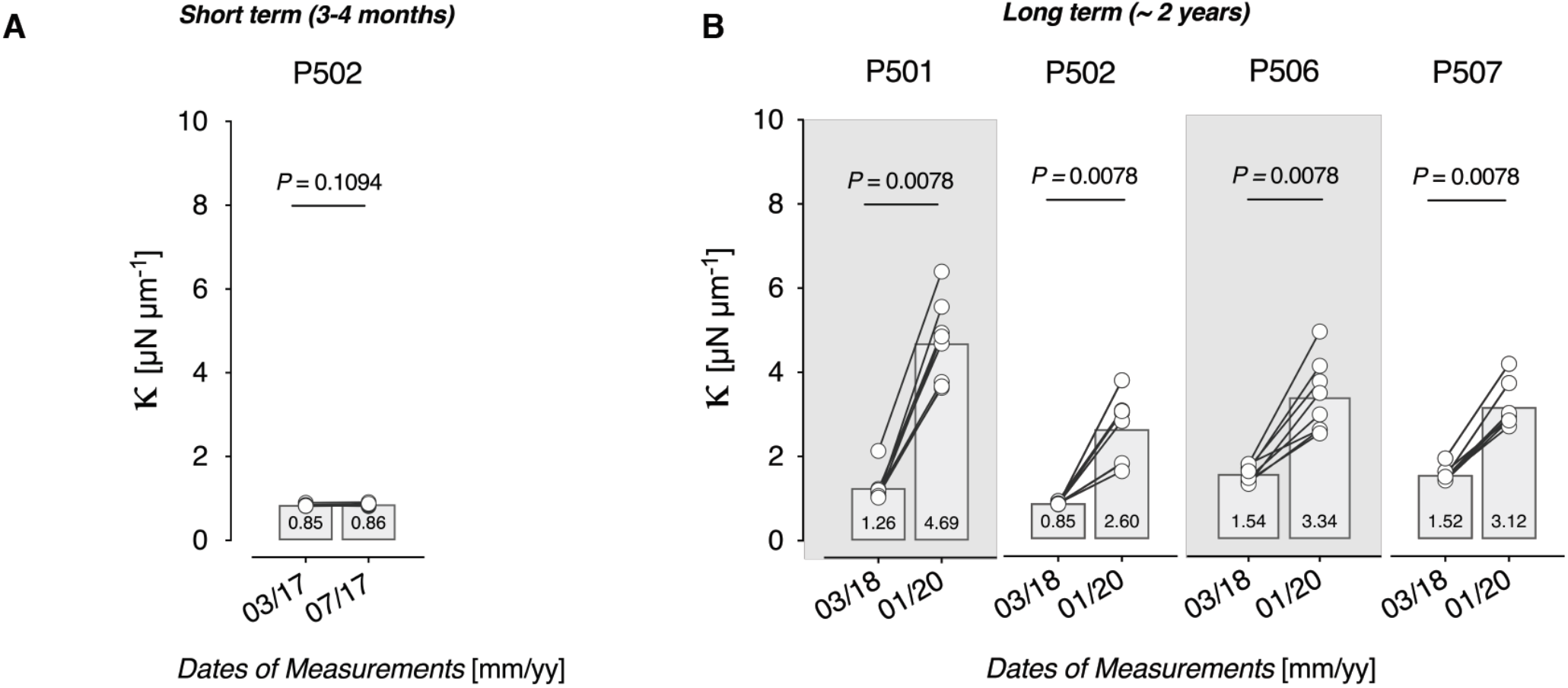
Evaluation of the short- and long-term evolution of the steel posts spring constant (*k*) A well-known shortcoming of Sylgard 184 PDMS is that it contains high amounts of silica impurities which interfere with heat curing, especially when mixed in non-stoichiometric ratios (*e.g.* 50:1 or 100:1). We addressed this limitation by re-calibrating the posts spring constant at a 3 to 4-month interval. (*n* = 8 posts/plate, Nonparametric, Wilcoxon matched-pairs signed rank test.

**Supplementary figure 2.**
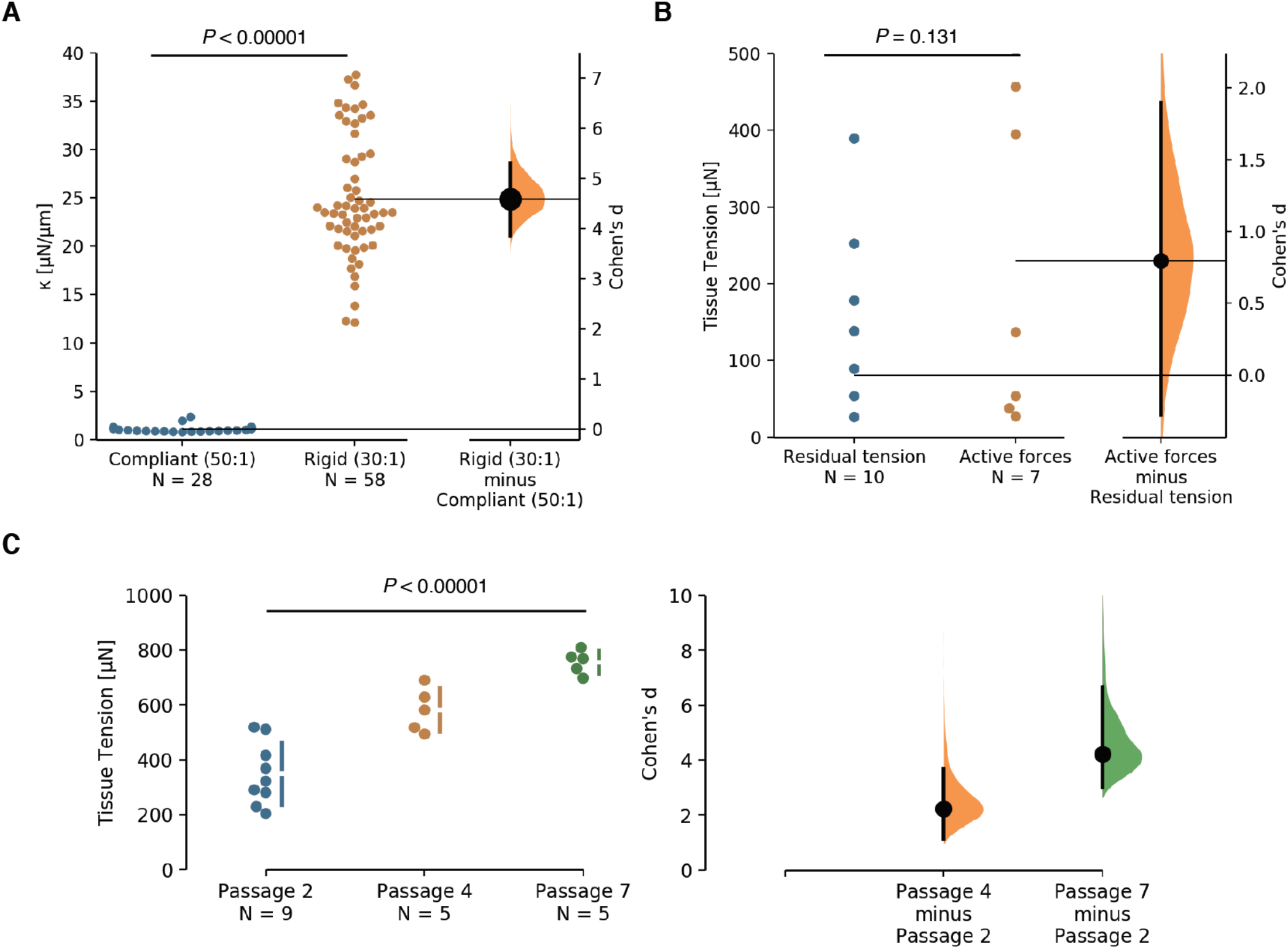
Gardner–Altman plots for the estimation statistics for data in Figure 1. The difference axis of the estimation plot displays the effect size, here the Cohen’s *d*. The effect sizes and CIs are reported above as: effect size [CI width lower bound; upper bound]. The 95% confidence interval of Cohen’s *d* is illustrated by the black vertical line. The curve displays the distribution of 5000 bootstrap re-samplings. *P* value denotes the two-sided permutation. **(A)** Post spring constant (*k*). The unpaired Cohen’s d between Compliant (50:1) and Rigid (30:1) is 4.59 [95.0%CI 3.85, 5.3]. **(B)** Active and residual tension at the end of the 48h timepoint of vehicle vs stimulants conditions (*n* = 7-10 tissues). The unpaired Cohen’s d between Residual tension and Active forces is 0.797 [95.0%CI −0.278, 1.9]. **(C)** Quantification of tissue tension as a function of the passage number of rat tail-derived tendon stromal cells (*n* = 5-9 tissues). The unpaired Cohen’s d between Passage 2 and Passage 4 is 2.24 [95.0%CI 1.13, 3.67]. The unpaired Cohen’s d between Passage 2 and Passage 7 is 4.23 [95.0%CI 3.0, 6.68].

**Supplementary figure 3.**
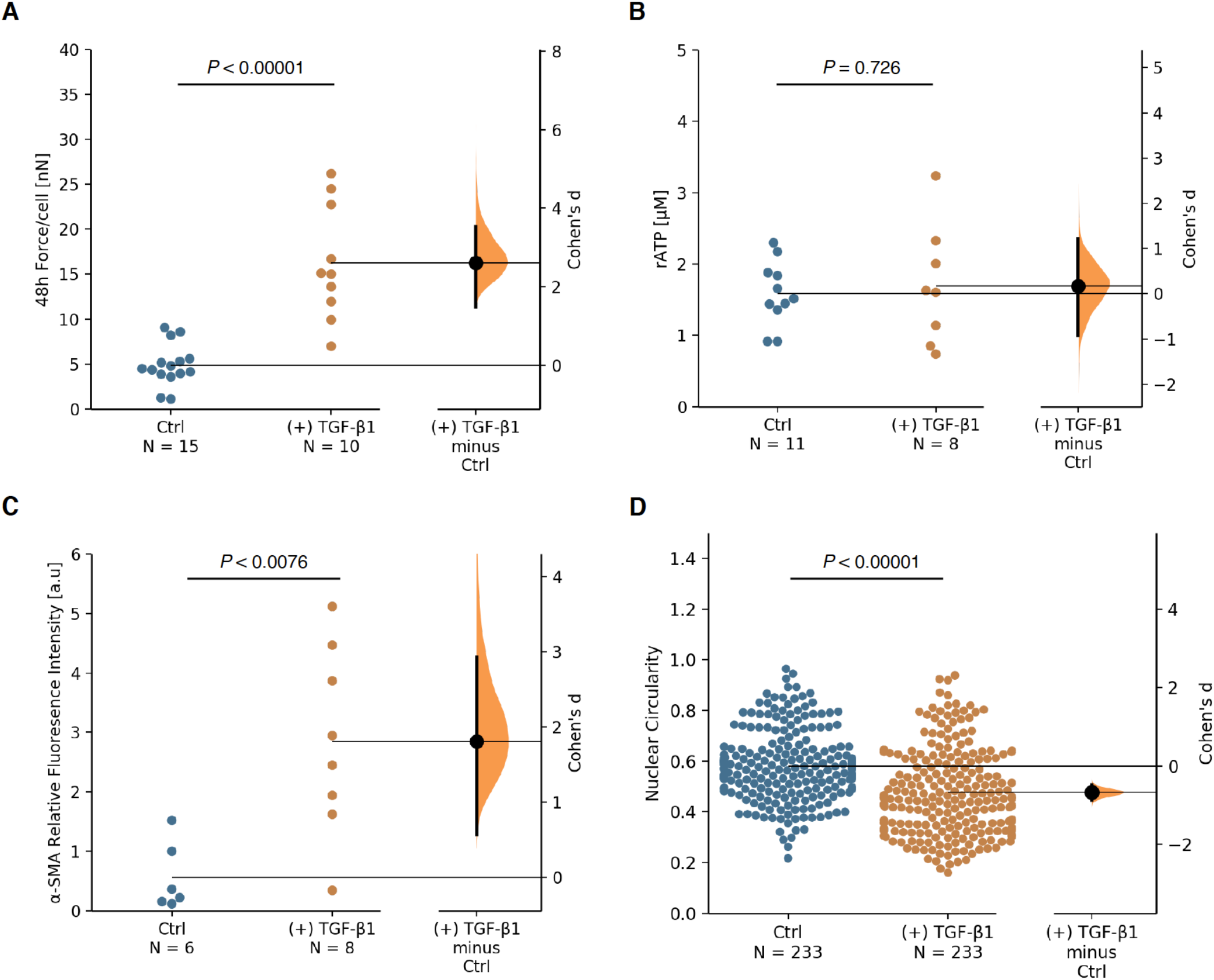
Gardner–Altman plots for the estimation statistics for data in Figure 2. The difference axis of the estimation plot displays the effect size, here the Cohen’s *d*. The effect sizes and CIs are reported above as: effect size [CI width lower bound; upper bound]. The 95% confidence interval of Cohen’s *d* is illustrated by the black vertical line. The curve displays the distribution of 5000 bootstrap re-samplings. *P* value denotes the two-sided permutation. **(A)** Quantitative analysis of tissue traction forces, as force per cell. (*n* = 10-15 tissues). The unpaired Cohen’s d between Ctrl and (+) TGF-β1 is 2.61 [95.0%CI 1.48, 3.52]. **(B)** Analysis of cellular metabolic activity, as a measure of viability. (*n* = 8-11 tissues). The unpaired Cohen’s d between Ctrl and (+) TGF-β1 is 0.168 [95.0%CI −0.924, 1.21]. **(C)** Quantification of fluorescent intensity levels of smooth muscle alpha-actin (α-SMA) in TGF-β1 stimulated and vehicle control. (*n* = 6-8 tissues from 2 biologically independent experiments). The unpaired Cohen’s d between Ctrl and (+) TGF-β1 is 1.8 [95.0%CI 0.57, 2.93]. **(D)** Quantification of nuclear circularity of tendon-derived stromal cells. The unpaired Cohen’s d between Ctrl and (+) TGF-β1 is −0.674 [95.0%CI −0.87, −0.48].

**Supplementary figure 4.**
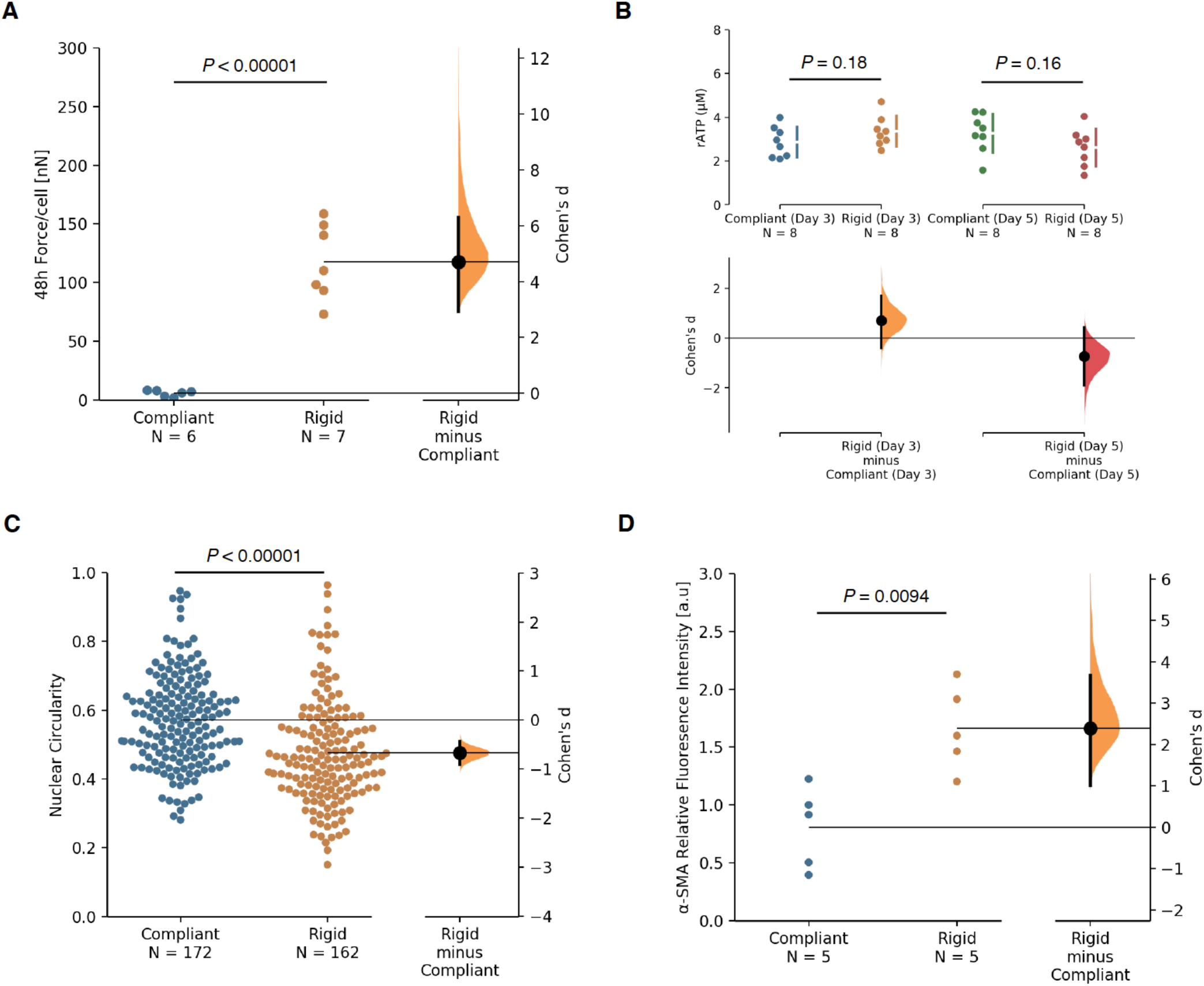
Gardner–Altman plots for the estimation statistics for data in Figure 3. The difference axis of the estimation plot displays the effect size, here the Cohen’s *d*. The effect sizes and CIs are reported above as: effect size [CI width lower bound; upper bound]. The 95% confidence interval of Cohen’s *d* is illustrated by the black vertical line. The curve displays the distribution of 5000 bootstrap re-samplings. *P* value denotes the two-sided permutation. **(A)** Quantitative analysis of tissue traction forces, as force per cell (*n* = 6-7 tissues). The unpaired Cohen’s d between Compliant and Rigid is 4.7 [95.0%CI 2.93, 6.31]. **(B)** Analysis of cellular metabolic activity, as a measure of viability. (*n* = 8 tissues/group from 2 biologically independent experiments). The unpaired Cohen’s d between Compliant_ Day3 and Rigid_ Day3 is 0.692 [95.0%CI −0.396, 1.7]. The unpaired Cohen’s d between Compliant_ Day5 and Rigid_ Day5 is −0.74 [95.0%CI −1.91, 0.413]. **(C)** Quantification of nuclear circularity of tendon-derived stromal cells. The unpaired Cohen’s d between Compliant and Rigid is −0.671 [95.0%CI −0.903, −0.442]. **(D)** Quantification of fluorescent intensity levels of smooth muscle alpha-actin (α-SMA), as a function of boundary stiffness. (*n* = 5 tissues from 2 biologically independent experiments). The unpaired Cohen’s d between Compliant and Rigid is 2.39 [95.0%CI 0.999, 3.67].

**Supplementary figure 5.**
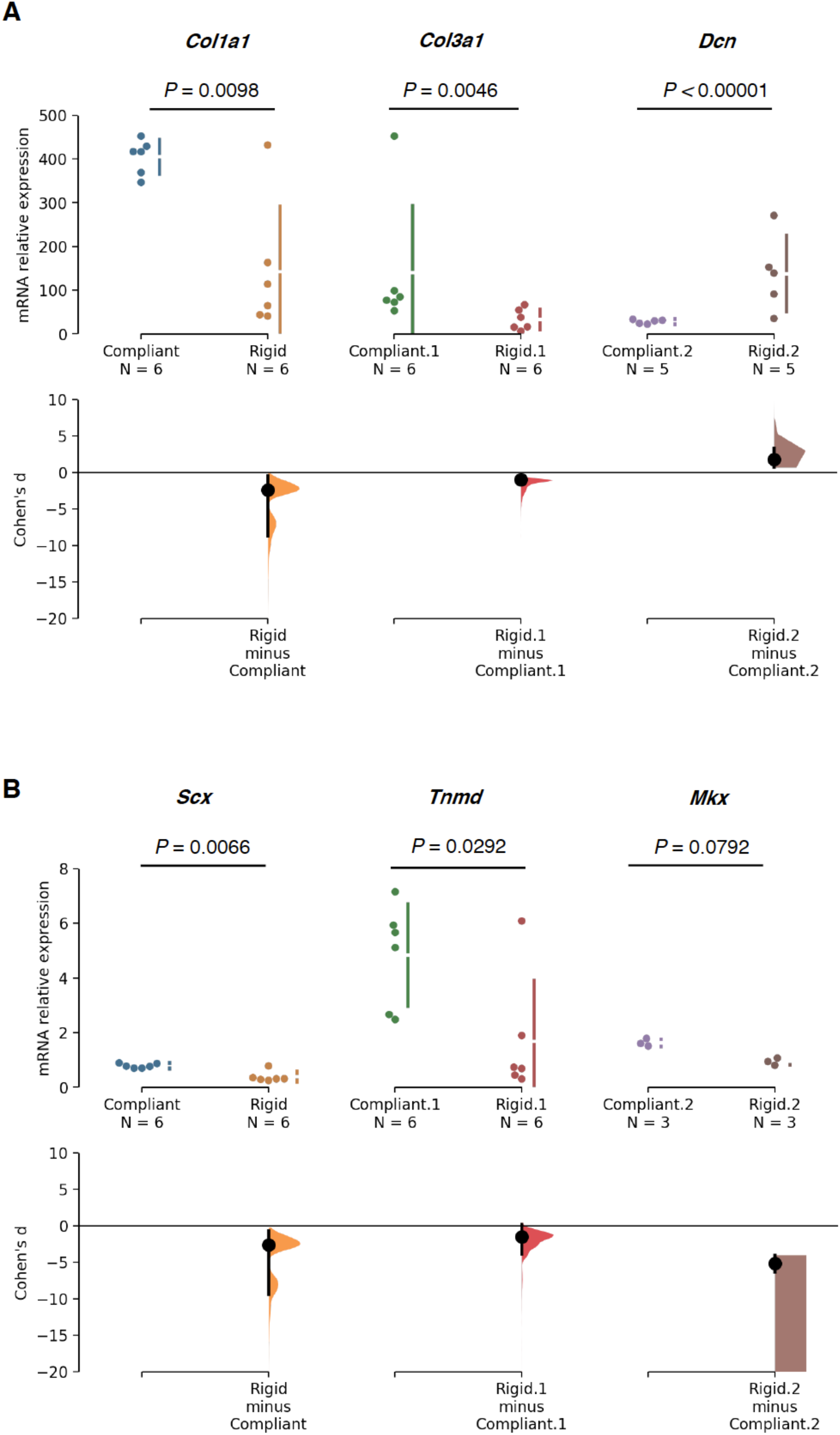
Gardner–Altman plots for the estimation statistics for data in Figure 3. The difference axis of the estimation plot displays the effect size, here the Cohen’s *d*. The effect sizes and CIs are reported above as: effect size [CI width lower bound; upper bound]. The 95% confidence interval of Cohen’s *d* is illustrated by the black vertical line. The curve displays the distribution of 5000 bootstrap re-samplings. *P* value denotes the two-sided permutation. **(A)** mRNA expression of ECM-related genes, and **(B)** tendon lineage-related genes in stromal cells tethered to different mechanical rigidities. (*n* = 3-6 replicates/group from 3 biologically independent experiments, with each data point representing a ΔCt value of 2-3 pooled tissues. The unpaired Cohen’s d between Col1a1 Compliant and Rigid is −2.4 [95.0%CI −8.76, −0.442]. The unpaired Cohen’s d between Col3a1 Compliant.1 and Rigid.1 is −0.969 [95.0%CI −1.28, −0.597]. The unpaired Cohen’s d between Dcn Compliant.2 and Rigid.2 is 1.77 [95.0%CI 0.716, 3.32]. The unpaired Cohen’s d between Scx Compliant and Rigid is −2.66 [95.0%CI −9.4, −0.706]. The unpaired Cohen’s d between Tnmd Compliant.1 and Rigid.1 is −1.53 [95.0%CI −3.87, 0.204]. The unpaired Cohen’s d between Mkx Compliant.2 and Rigid.2 is −5.14 [95.0%CI −6.32, −4.02].

**Supplementary figure 6.**
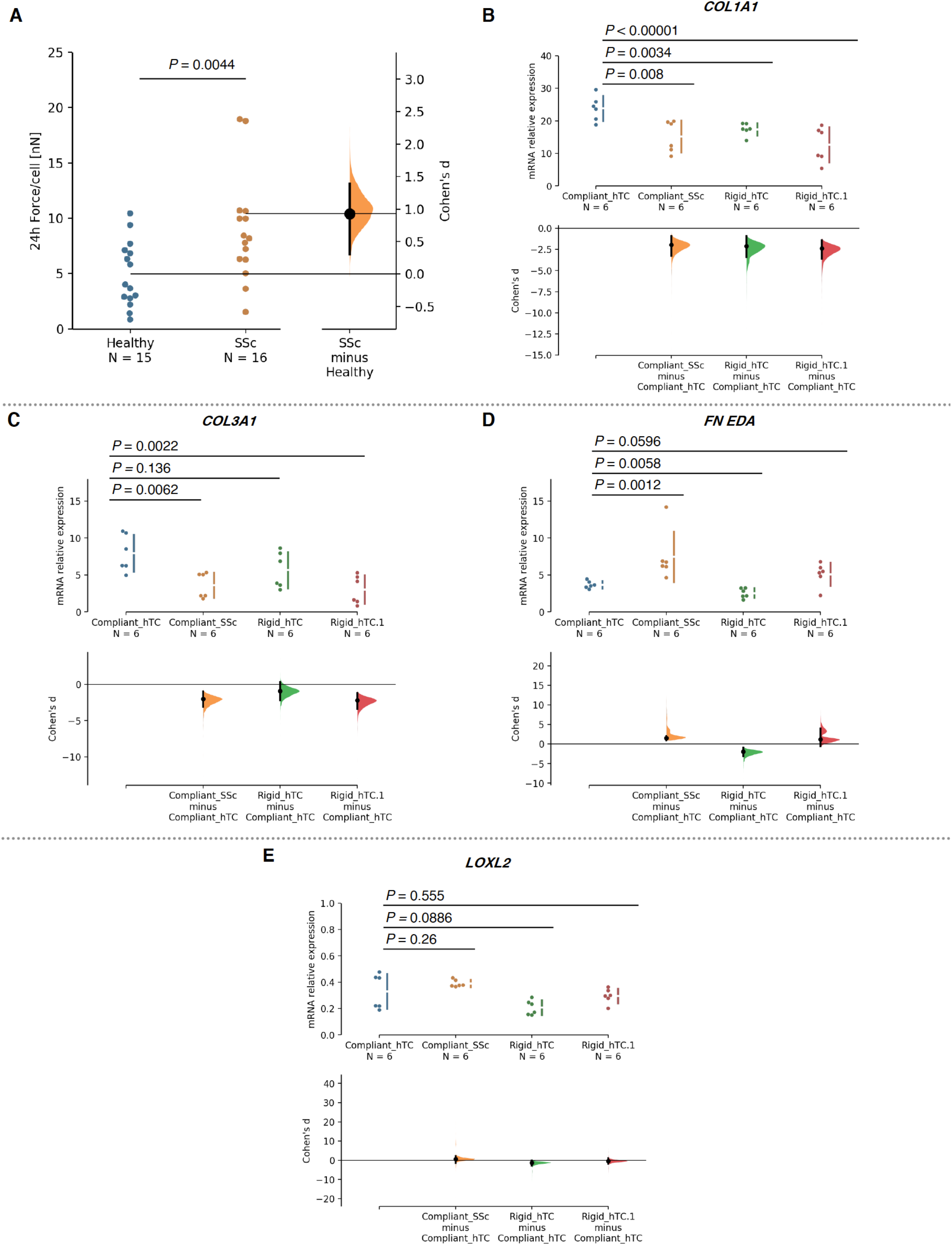
Gardner–Altman plots for the estimation statistics for data in Figure 4. The difference axis of the estimation plot displays the effect size, here the Cohen’s *d*. The effect sizes and CIs are reported above as: effect size [CI width lower bound; upper bound]. The 95% confidence interval of Cohen’s *d* is illustrated by the black vertical line. The curve displays the distribution of 5000 bootstrap re-samplings. *P* value denotes the two-sided permutation. **(A)** Quantitative analysis of tissue traction forces, as force per cell (*n* = 24 tissues/group from 3 anatomically different tendons of the SSc donor and their matching controls from 3 independent healthy donors). The unpaired Cohen’s d between Healthy and SSc is 0.928 [95.0%CI 0.305, 1.39]. mRNA expression of ECM-related genes of systemic sclerosis (SSc) and healthy (hTC) tendon-derived stromal cells tethered to different mechanical rigidities. **(B) *COL1A1*.** The unpaired Cohen’s d between Compliant_hTC and Compliant_SSc is −1.96 [95.0%CI −3.27, −0.915]. The unpaired Cohen’s d between Compliant_hTC and Rigid_hTC is −2.11 [95.0%CI −3.42, −0.936]. The unpaired Cohen’s d between Compliant_hTC and Rigid_SSc is −2.38 [95.0%CI −3.61, −1.4]. **(C) *COL3A1*.** The unpaired Cohen’s d between Compliant_hTC and Compliant_SSc is −2.02 [95.0%CI −3.09, −0.963]. The unpaired Cohen’s d between Compliant_hTC and Rigid_hTC is −0.922 [95.0%CI −2.21, 0.302]. The unpaired Cohen’s d between Compliant_hTC and Rigid_hTC.1 is −2.21 [95.0%CI −3.39, −1.16]. **(C) *FN^(EDA)^*.** The unpaired Cohen’s d between Compliant_hTC and Compliant_SSc is 1.56 [95.0%CI 0.946, 2.06]. The unpaired Cohen’s d between Compliant_hTC and Rigid_hTC is −1.97 [95.0%CI −3.13, −0.875]. The unpaired Cohen’s d between Compliant_hTC and Rigid_hTC.1 is 1.23 [95.0%CI −0.587, 4.03]. **(D) *LOXL2*.** The unpaired Cohen’s d between Compliant_hTC and Compliant_SSc is 0.637 [95.0%CI −1.47, 2.16]. The unpaired Cohen’s d between Compliant_hTC and Rigid_hTC is −1.2 [95.0%CI −2.75, −0.084]. The unpaired Cohen’s d between Compliant_hTC and Rigid_hTC.1 is −0.331 [95.0%CI −1.72, 1.13].

**Supplementary figure 7.**
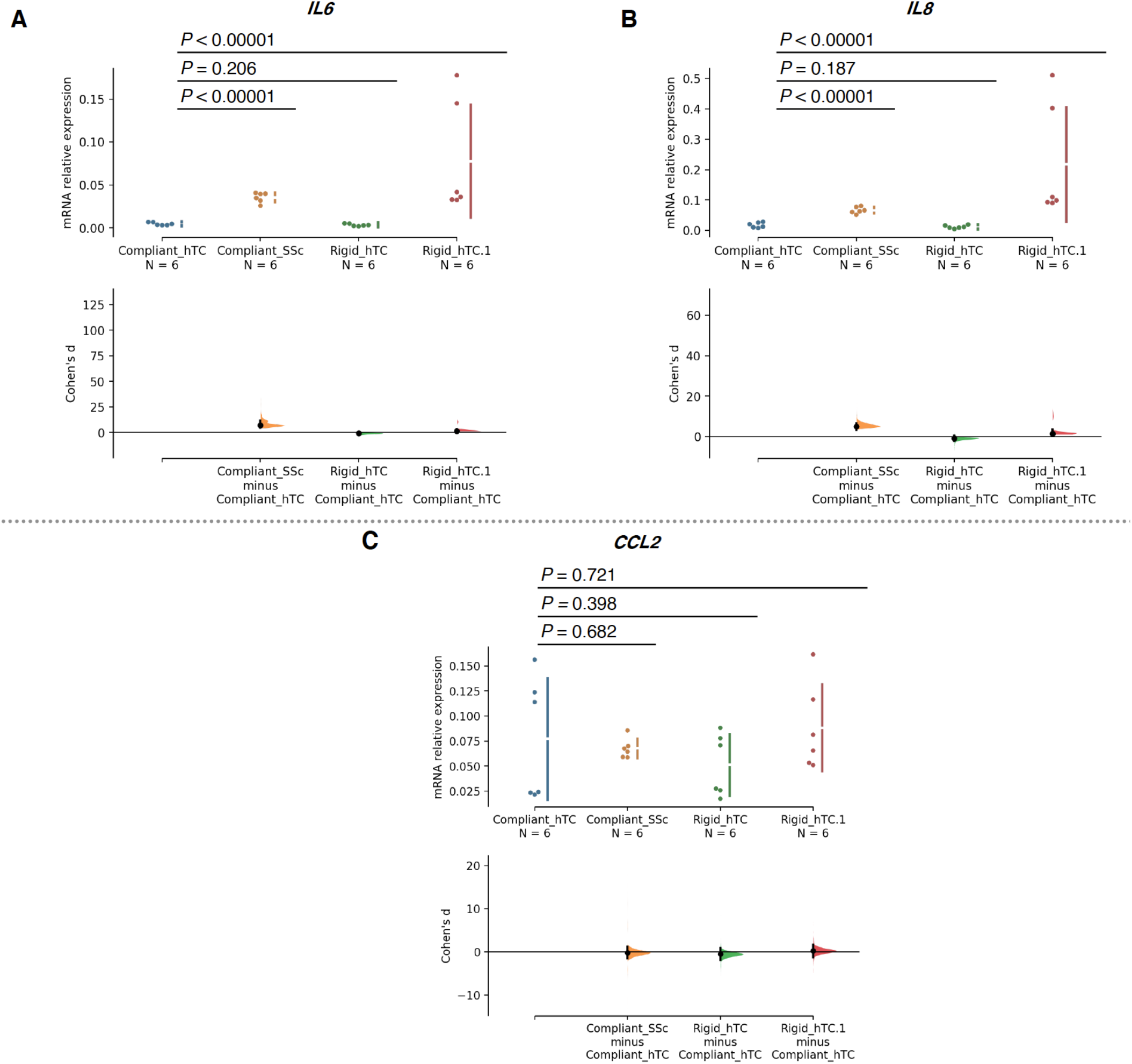
Gardner–Altman plots for the estimation statistics for data in Figure 4. The difference axis of the estimation plot displays the effect size, here the Cohen’s *d*. The effect sizes and CIs are reported above as: effect size [CI width lower bound; upper bound]. The 95% confidence interval of Cohen’s *d* is illustrated by the black vertical line. The curve displays the distribution of 5000 bootstrap re-samplings. *P* value denotes the two-sided permutation. mRNA expression of immune/inflammatory genes of systemic sclerosis (SSc) and healthy (hTC) tendon-derived stromal cells tethered to different mechanical rigidities. **(A) *IL6*.** The unpaired Cohen’s d between Compliant_hTC and Compliant_SSc is 7.12 [95.0%CI 4.56, 11.9]. The unpaired Cohen’s d between Compliant_hTC and Rigid_hTC is −0.784 [95.0%CI −2.1, 0.487]. The unpaired Cohen’s d between Compliant_hTC and Rigid_hTC.1 is 1.57 [95.0%CI 1.25, 3.58]. **(B) *IL8*.** The unpaired Cohen’s d between Compliant_hTC and Compliant_SSc is 5.03 [95.0%CI 3.39, 6.55]. The unpaired Cohen’s d between Compliant_hTC and Rigid_hTC is −0.824 [95.0%CI −2.14, 0.443]. The unpaired Cohen’s d between Compliant_hTC and Rigid_hTC.1 is 1.5 [95.0%CI 1.17, 3.57]. **(C) *CCL2*.** The unpaired Cohen’s d between Compliant_hTC and Compliant_SSc is −0.22 [95.0%CI −1.55, 1.17]. The unpaired Cohen’s d between Compliant_hTC and Rigid_hTC is −0.536 [95.0%CI −1.93, 0.876]. The unpaired Cohen’s d between Compliant_hTC and Rigid_hTC.1 is 0.209 [95.0%CI −1.2, 1.66].

**Supplementary figure 8.**
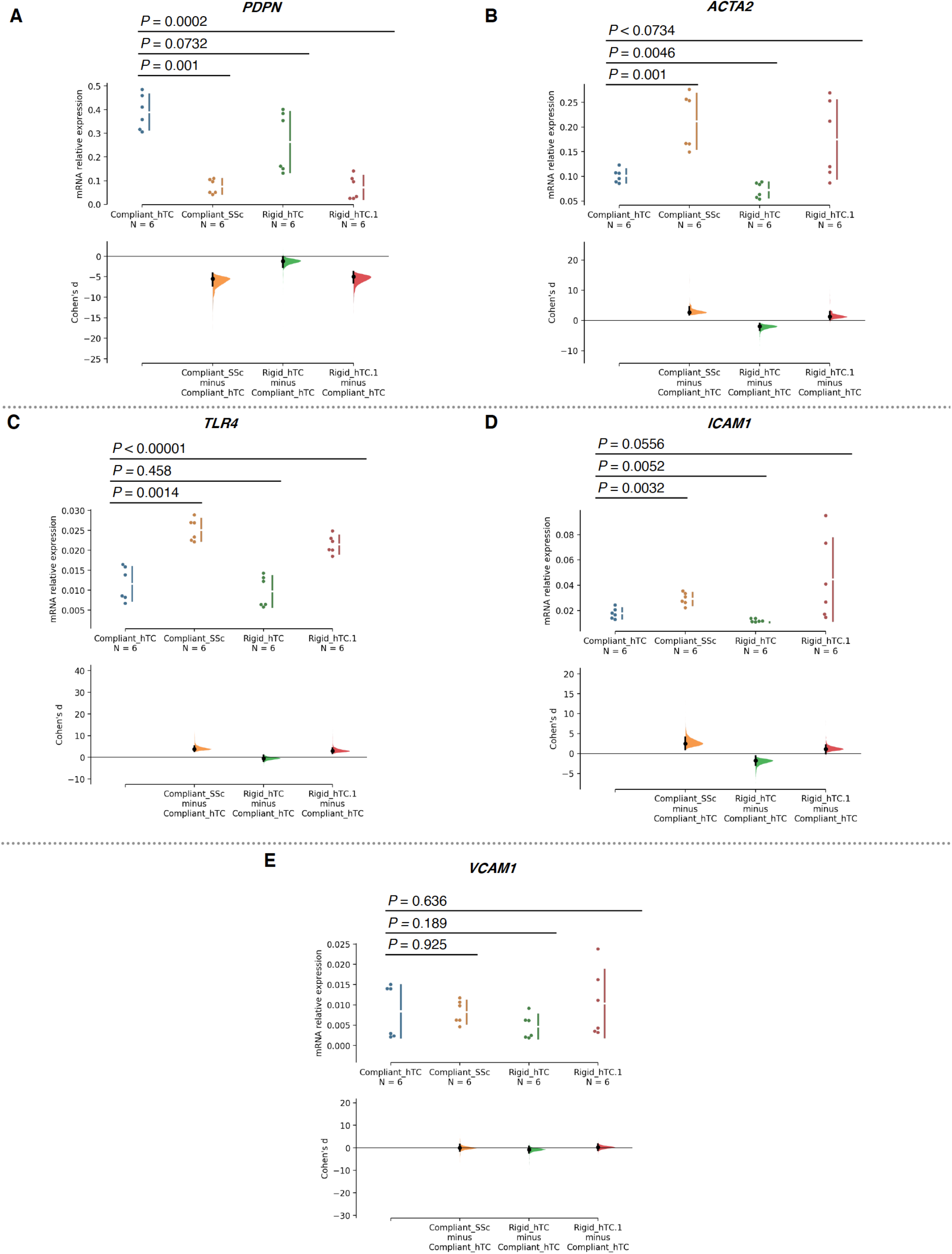
Gardner–Altman plots for the estimation statistics for data in Figure 4. The difference axis of the estimation plot displays the effect size, here the Cohen’s *d*. The effect sizes and CIs are reported above as: effect size [CI width lower bound; upper bound]. The 95% confidence interval of Cohen’s *d* is illustrated by the black vertical line. The curve displays the distribution of 5000 bootstrap re-samplings. *P* value denotes the two-sided permutation. mRNA expression of stromal activation markers genes of systemic sclerosis (SSc) and healthy (hTC) tendon-derived stromal cells tethered to different mechanical rigidities. **(A) *PDPN*.** The unpaired Cohen’s d between Compliant_hTC and Compliant_SSc is −5.49 [95.0%CI −7.2, −4.17]. The unpaired Cohen’s d between Compliant_hTC and Rigid_hTC is −1.2 [95.0%CI −2.69, −0.0865]. The unpaired Cohen’s d between Compliant_hTC and Rigid_hTC.1 is −5.0 [95.0%CI −6.44, −3.75]. **(B) *ACTA2*.** The unpaired Cohen’s d between Compliant_hTC and Compliant_SSc is 2.69 [95.0%CI 1.9, 4.42]. The unpaired Cohen’s d between Compliant_hTC and Rigid_hTC is −1.95 [95.0%CI −3.22, −0.998]. The unpaired Cohen’s d between Compliant_hTC and Rigid_hTC.1 is 1.29 [95.0%CI 0.164, 2.88]. **(C) *TLR4*.** The unpaired Cohen’s d between Compliant_hTC and Compliant_SSc is 3.74 [95.0%CI 2.73, 5.08]. The unpaired Cohen’s d between Compliant_hTC and Rigid_hTC is −0.464 [95.0%CI −1.78, 0.805]. The unpaired Cohen’s d between Compliant_hTC and Rigid_hTC.1 is 2.87 [95.0%CI 1.87, 4.21]. **(D) *ICAM1*.** The unpaired Cohen’s d between Compliant_hTC and Compliant_SSc is 2.52 [95.0%CI 1.04, 4.09]. The unpaired Cohen’s d between Compliant_hTC and Rigid_hTC is −1.77 [95.0%CI −2.88, −0.631]. The unpaired Cohen’s d between Compliant_hTC and Rigid_hTC.1 is 1.15 [95.0%CI 0.0216, 2.11]. **(E) *VCAM1*.** The unpaired Cohen’s d between Compliant_hTC and Compliant_SSc is −0.0376 [95.0%CI −1.46, 1.49]. The unpaired Cohen’s d between Compliant_hTC and Rigid_hTC is −0.739 [95.0%CI −2.23, 0.617]. The unpaired Cohen’s d between Compliant_hTC and Rigid_hTC.1 is 0.26 [95.0%CI −1.06, 1.63].

**Supplementary figure 9.**
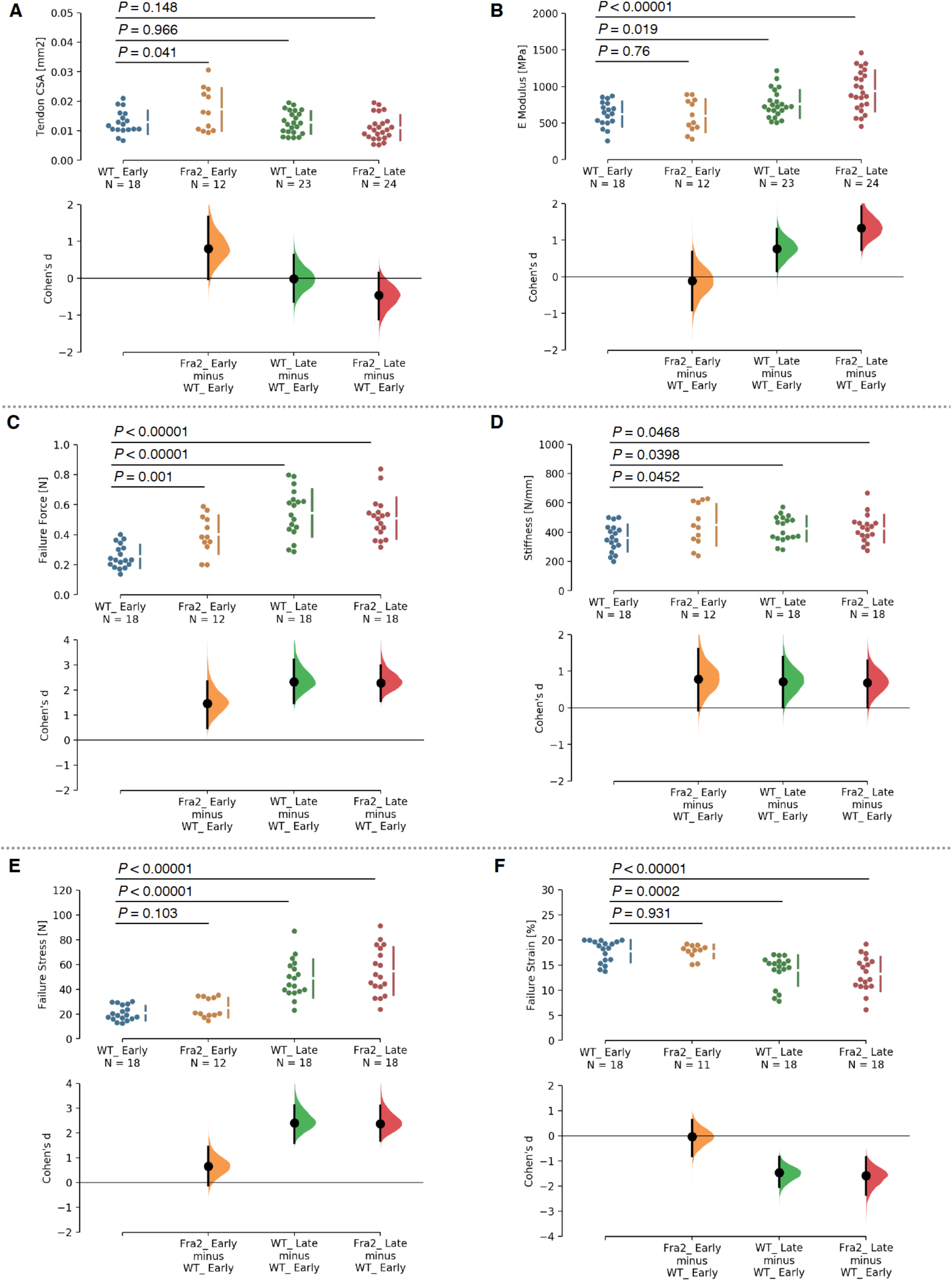
Gardner–Altman plots for the estimation statistics for data in Figure 5. The difference axis of the estimation plot displays the effect size, here the Cohen’s *d*. The effect sizes and CIs are reported above as: effect size [CI width lower bound; upper bound]. The 95% confidence interval of Cohen’s *d* is illustrated by the black vertical line. The curve displays the distribution of 5000 bootstrap re-samplings. *P* value denotes the two-sided permutation. Biomechanical characterization of Fra2^Tg^ tendons and their WT littermates at early vs. late-stages of established fibrosis. Each data point represents an independent sample; (*n* = 12-24 fascicles from 6-7 mice/group). **(A) Tail tendon cross-sectional area (CSA)**. The unpaired Cohen’s d between WT_ Early and Fra2_ Early is 0.8 [95.0%CI −0.0207, 1.67]. The unpaired Cohen’s d between WT_ Early and WT_ Late is −0.0135 [95.0%CI −0.642, 0.633]. The unpaired Cohen’s d between WT_ Early and Fra2_ Late is −0.461 [95.0%CI −1.12, 0.152]. **(B) *E* modulus**. The unpaired Cohen’s d between WT_ Early and Fra2_ Early is −0.113 [95.0%CI −0.913, 0.681]. The unpaired Cohen’s d between WT_ Early and WT_ Late is 0.763 [95.0%CI 0.145, 1.31]. The unpaired Cohen’s d between WT_ Early and Fra2_ Late is 1.33 [95.0%CI 0.738, 1.92]. **(C) Failure force**. The unpaired Cohen’s d between WT_ Early and Fra2_ Early is 1.46 [95.0%CI 0.47, 2.34]. The unpaired Cohen’s d between WT_ Early and WT_ Late is 2.33 [95.0%CI 1.48, 3.21]. The unpaired Cohen’s d between WT_ Early and Fra2_ Late is 2.29 [95.0%CI 1.57, 2.98]. **(D) Stiffness**. The unpaired Cohen’s d between WT_ Early and Fra2_ Early is 0.784 [95.0%CI −0.0689, 1.61]. The unpaired Cohen’s d between WT_ Early and WT_ Late is 0.718 [95.0%CI 0.0101, 1.39]. The unpaired Cohen’s d between WT_ Early and Fra2_ Late is 0.688 [95.0%CI 0.00584, 1.3]. **(E) Failure stress**. The unpaired Cohen’s d between WT_ Early and Fra2_ Early is 0.649 [95.0%CI −0.115, 1.45]. The unpaired Cohen’s d between WT_ Early and WT_ Late is 2.4 [95.0%CI 1.59, 3.12]. The unpaired Cohen’s d between WT_ Early and Fra2_ Late is 2.38 [95.0%CI 1.69, 3.11]. **(F) Failure strain**. The unpaired Cohen’s d between WT_ Early and Fra2_ Early is −0.0332 [95.0%CI −0.8, 0.63]. The unpaired Cohen’s d between WT_ Early and WT_ Late is −1.46 [95.0%CI −2.04, −0.834]. The unpaired Cohen’s d between WT_ Early and Fra2_ Late is −1.58 [95.0%CI −2.34, −0.858].

**Supplementary figure 10.**
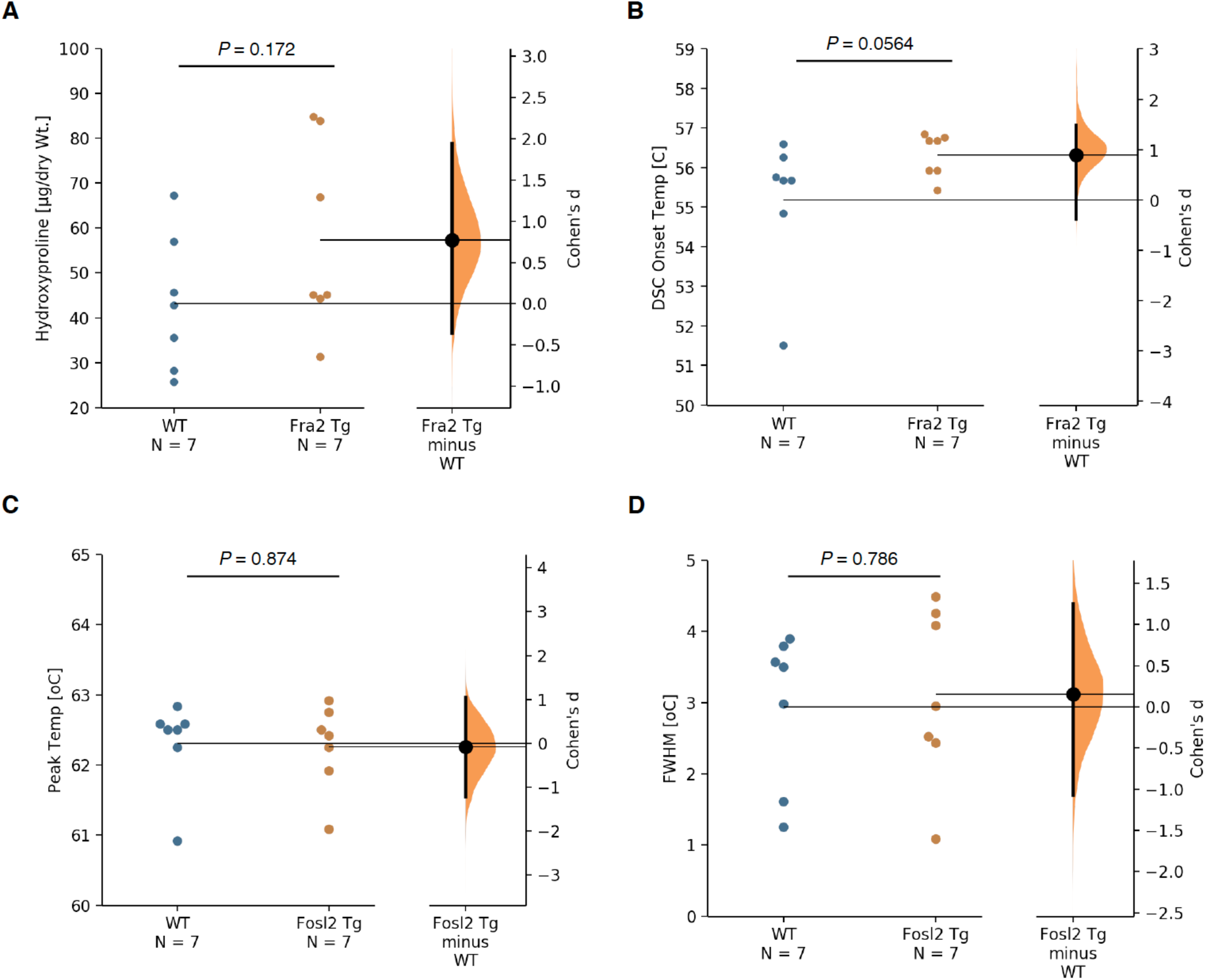
Gardner–Altman plots for the estimation statistics for data in Figure 5. The difference axis of the estimation plot displays the effect size, here the Cohen’s *d*. The effect sizes and CIs are reported above as: effect size [CI width lower bound; upper bound]. The 95% confidence interval of Cohen’s *d* is illustrated by the black vertical line. The curve displays the distribution of 5000 bootstrap re-samplings. *P* value denotes the two-sided permutation. **(A)** Quantification of hydroxyproline content in tendons (*n* = 7 mice/genotype). The unpaired Cohen’s d between WT and Fra2 Tg is 0.771 [95.0%CI −0.353, 1.94]. Thermal denaturing of tendons as measured by DSC (*n* = 7 mice/genotype). **(B)** DSC Endothermic onset temperature (°C).The unpaired Cohen’s d between WT and Fosl2 Tg is 0.889 [95.0%CI −0.376, 1.48]. **(C)** Peak temperature (°C).The unpaired Cohen’s d between WT and Fosl2 Tg is −0.0761 [95.0%CI −1.22, 1.04]. **(D)** Full-width at half-maximum (FWHM). The unpaired Cohen’s d between WT and Fosl2 Tg is 0.15 [95.0%CI −1.07, 1.26].

**Supplementary figure 11.**
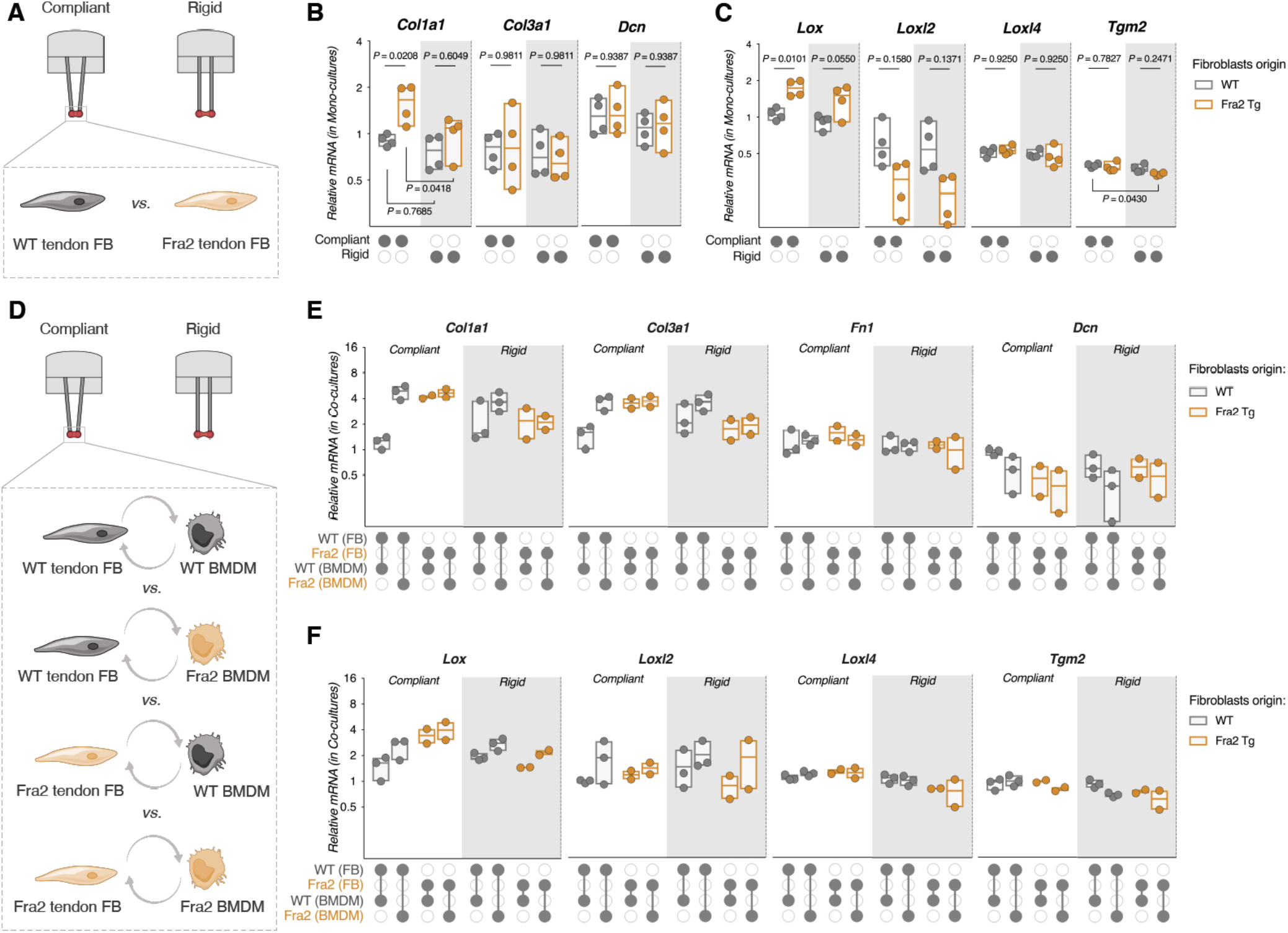
Macrophages enhance stromal fibroblasts activation and collagen expression, but not ECM crosslinking enzymes. **(A)** Experimental design for monocultures. **(B)** mRNA expression of ECM-related genes, **(C)** ECM crosslinking enzymes in Fra-2 Tg and WT cells tethered to different mechanical rigidities. (*n* = 4 mice/genotype. Each data point represents fold-change value, horizontal lines indicate the median, Two-way ANOVA (boundary stiffness, genotype) with Tukey’s *post-hoc* test). **(D)** Experimental design for “mix-and-match” direct co-cultures. **(E)** mRNA expression of ECM-related genes, **(F)** ECM crosslinking enzymes in Fra-2 Tg and WT cells tethered to different mechanical rigidities. (WT: *n* = 4 mice – Fra-2 Tg: *n* = 2 mice). Each data point represents fold-change value, horizontal lines indicate the median, Two-way ANOVA (boundary stiffness, genotype) with Tukey’s *post-hoc* test). Estimation plots and permuted *P* values for (B and C) are in (Supplementary figure 12 and 13). Cohen’s *d* effect sizes and CIs are reported above as: Cohen’s *d* [CI width lower bound; upper bound].

**Supplementary figure 12.**
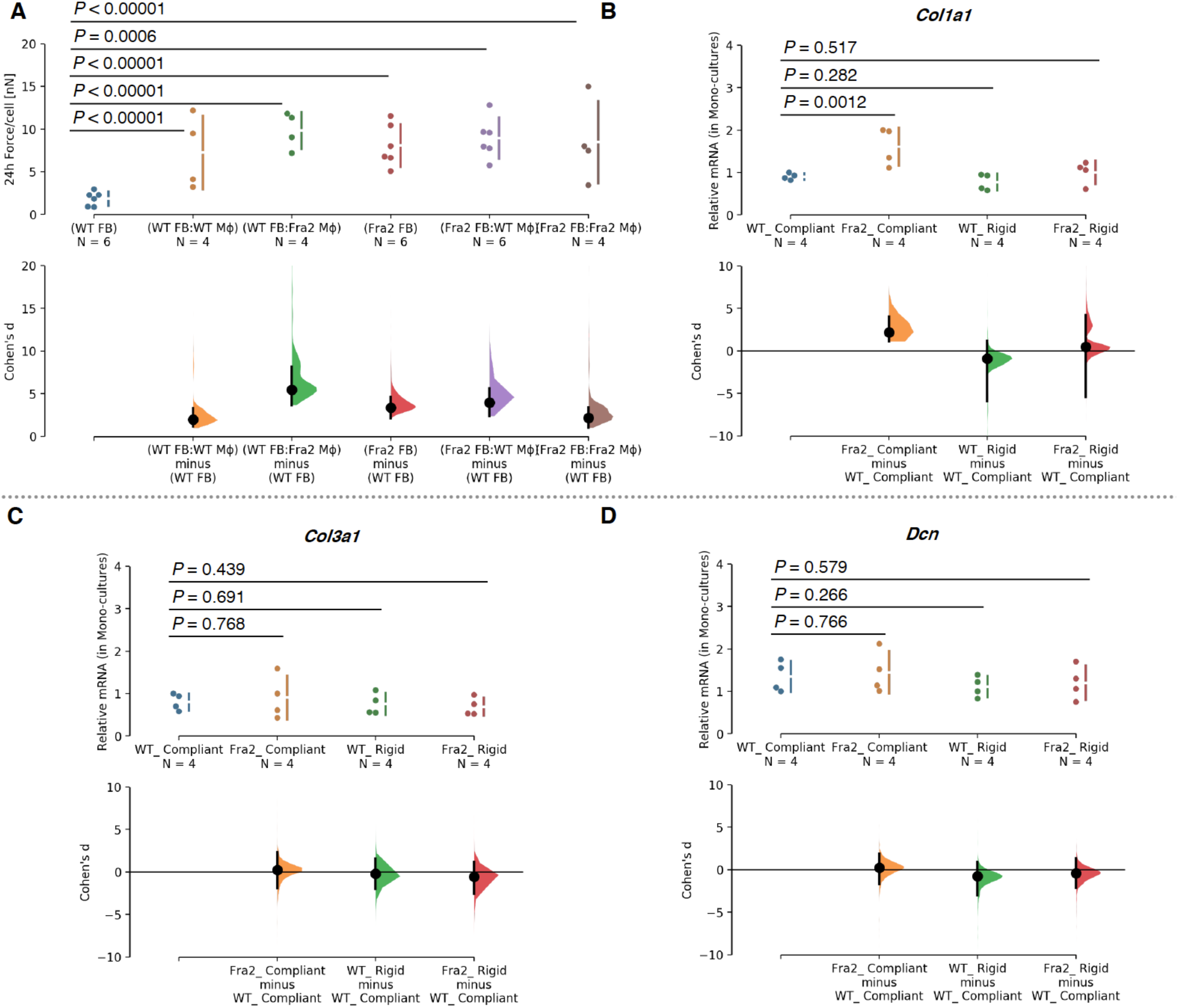
Gardner–Altman plots for the estimation statistics for data in Figure 6 and Supplementary figure 9. The difference axis of the estimation plot displays the effect size, here the Cohen’s *d*. The effect sizes and CIs are reported above as: effect size [CI width lower bound; upper bound]. The 95% confidence interval of Cohen’s *d* is illustrated by the black vertical line. The curve displays the distribution of 5000 bootstrap re-samplings. *P* value denotes the two-sided permutation. **(A)** Quantification of tissue traction forces per cell, following normalization to initial seeding density of stromal fibroblasts. *n* = 4-6 tissues from 2-4 mice/genotype. The unpaired Cohen’s d between (WT FB) and (WT FB:WT Mϕ) is 1.99 [95.0%CI 1.16, 3.34]. The unpaired Cohen’s d between (WT FB) and (WT FB:Fra2 Mϕ) is 5.44 [95.0%CI 3.68, 8.14]. The unpaired Cohen’s d between (WT FB) and (Fra2 FB) is 3.39 [95.0%CI 2.13, 4.59]. The unpaired Cohen’s d between (WT FB) and (Fra2 FB:WT Mϕ) is 3.95 [95.0%CI 2.39, 5.62]. The unpaired Cohen’s d between (WT FB) and (Fra2 FB:Fra2 Mϕ) is 2.2 [95.0%CI 1.05, 3.44]. mRNA expression of ECM-related genes of in Fra2 Tg and WT stromal cells tethered to different mechanical rigidities. **(B)** *Col1a1*. The unpaired Cohen’s d between WT_ Compliant and Fra2_ Compliant is 2.19 [95.0%CI 1.15, 4.04]. The unpaired Cohen’s d between WT_ Compliant and WT_ Rigid is −0.894 [95.0%CI −5.92, 1.17]. The unpaired Cohen’s d between WT_ Compliant and Fra2_ Rigid is 0.499 [95.0%CI −5.45, 4.2]. **(C)** *Col3a1*. The unpaired Cohen’s d between WT_ Compliant and Fra2_ Compliant is 0.263 [95.0%CI −1.88, 2.33]. The unpaired Cohen’s d between WT_ Compliant and WT_ Rigid is −0.209 [95.0%CI −2.0, 1.59]. The unpaired Cohen’s d between WT_ Compliant and Fra2_ Rigid is −0.546 [95.0%CI −2.52, 1.18]. **(D)** *Dcn*. The unpaired Cohen’s d between WT_ Compliant and Fra2_ Compliant is 0.23 [95.0%CI −1.72, 1.89]. The unpaired Cohen’s d between WT_ Compliant and WT_ Rigid is −0.77 [95.0%CI −2.99, 0.918]. The unpaired Cohen’s d between WT_ Compliant and Fra2_ Rigid is −0.38 [95.0%CI −2.16, 1.32].

**Supplementary figure 13.**
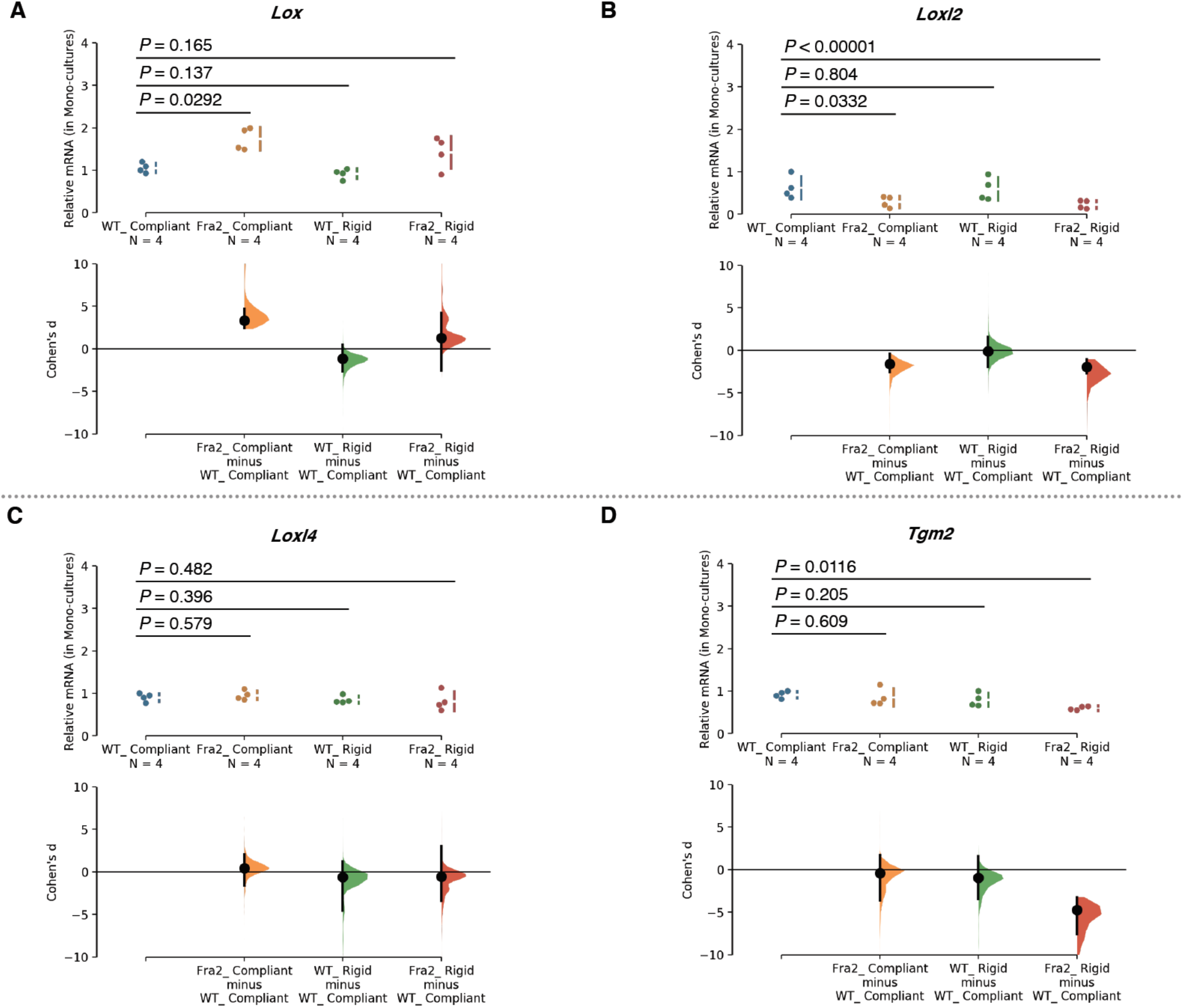
Gardner–Altman plots for the estimation statistics for data in Supplementary figure 9. The difference axis of the estimation plot displays the effect size, here the Cohen’s *d*. The effect sizes and CIs are reported above as: effect size [CI width lower bound; upper bound]. The 95% confidence interval of Cohen’s *d* is illustrated by the black vertical line. The curve displays the distribution of 5000 bootstrap re-samplings. *P* value denotes the two-sided permutation. mRNA expression of ECM-related genes of in Fra2 Tg and WT stromal cells tethered to different mechanical rigidities. mRNA expression of ECM-related genes of in Fra2 Tg and WT stromal cells tethered to different mechanical rigidities. **(A)** *Lox*. The unpaired Cohen’s d between WT_ Compliant and Fra2_ Compliant is 3.34 [95.0%CI 2.44, 4.73]. The unpaired Cohen’s d between WT_ Compliant and WT_ Rigid is −1.17 [95.0%CI −2.67, 0.491]. The unpaired Cohen’s d between WT_ Compliant and Fra2_ Rigid is 1.29 [95.0%CI −2.57, 4.23]. **(B)** *Loxl2*. The unpaired Cohen’s d between WT_ Compliant and Fra2_ Compliant is −1.59 [95.0%CI −2.55, −0.409]. The unpaired Cohen’s d between WT_ Compliant and WT_ Rigid is − 0.111 [95.0%CI −1.94, 1.61]. The unpaired Cohen’s d between WT_ Compliant and Fra2_ Rigid is −1.96 [95.0%CI −2.68, − 1.05]. **(C)** *Loxl4*. The unpaired Cohen’s d between WT_ Compliant and Fra2_ Compliant is 0.454 [95.0%CI −1.59, 2.1]. The unpaired Cohen’s d between WT_ Compliant and WT_ Rigid is −0.611 [95.0%CI −4.55, 1.23]. The unpaired Cohen’s d between WT_ Compliant and Fra2_ Rigid is −0.53 [95.0%CI −3.41, 3.03]. **(D)** *Tgm2*. The unpaired Cohen’s d between WT_ Compliant and Fra2_ Compliant is −0.413 [95.0%CI −3.62, 1.72]. The unpaired Cohen’s d between WT_ Compliant and WT_ Rigid is − 0.971 [95.0%CI −3.46, 1.57]. The unpaired Cohen’s d between WT_ Compliant and Fra2_ Rigid is −4.75 [95.0%CI −7.55, − 3.23].

**Supplementary Table S1:**
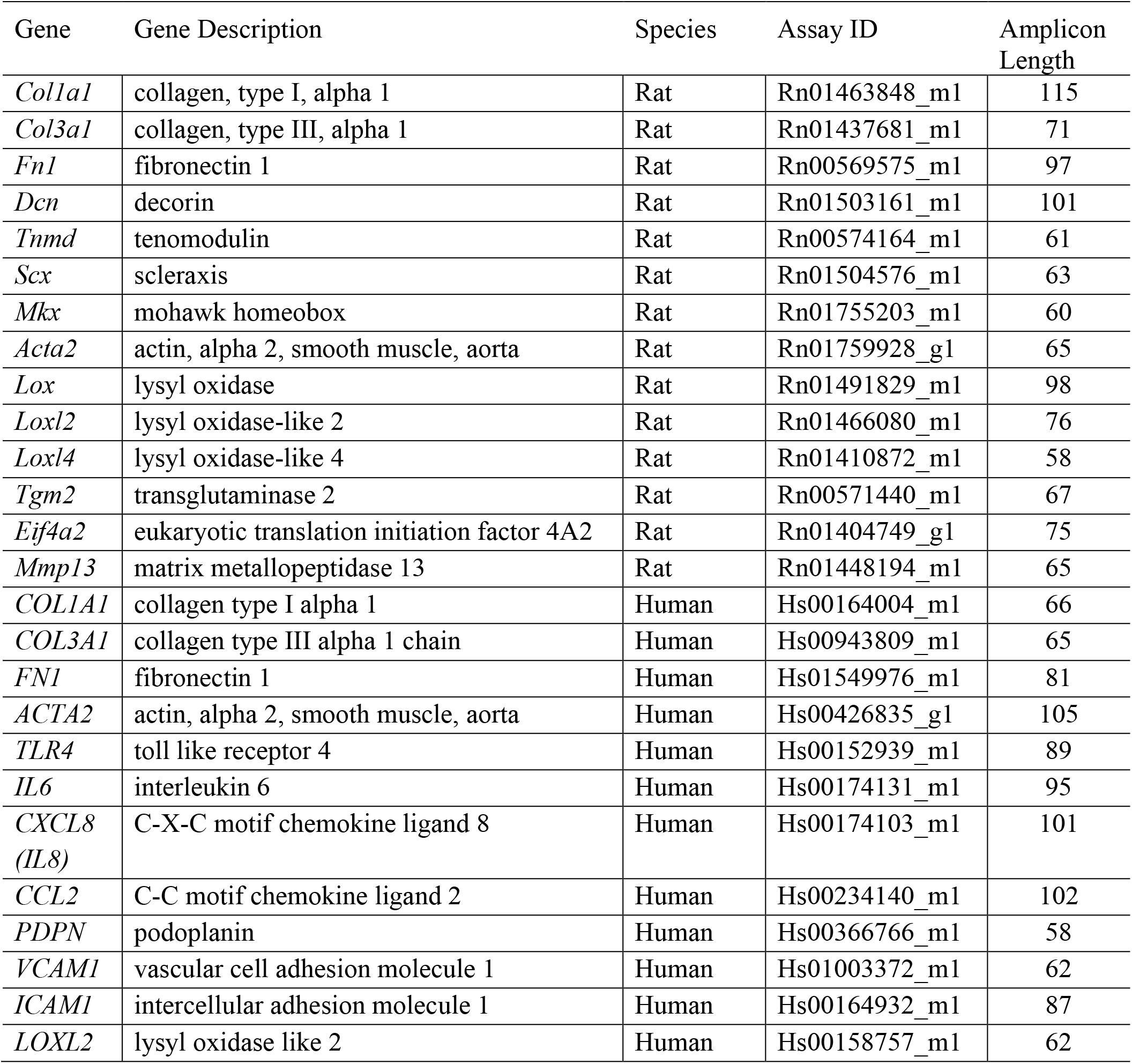
RT-qPCR TaqMan primers (Manufacturer: Thermo Fisher Scientific)

## References

1. Varga J, and Abraham D. Systemic sclerosis: a prototypic multisystem fibrotic disorder. J Clin Invest. 2007;117(3):557–67.

2. Gabrielli A, Avvedimento EV, and Krieg T. Scleroderma. N Engl J Med. 2009;360(19):1989–2003.

3. Allanore Y, Simms R, Distler O, Trojanowska M, Pope J, Denton CP, et al. Systemic sclerosis. Nature reviews Disease primers. 2015;1(1):15002.

4. Santos A, and Lagares D. Matrix Stiffness: the Conductor of Organ Fibrosis. Current Rheumatology Reports. 2018;20(1):2.

5. Disease GBD, Injury I, and Prevalence C. Global, regional, and national incidence, prevalence, and years lived with disability for 354 diseases and injuries for 195 countries and territories, 1990-2017: a systematic analysis for the Global Burden of Disease Study 2017. Lancet. 2018;392(10159):1789–858.

6. DALYs GBD, and Collaborators H. Global, regional, and national disability-adjusted life-years (DALYs) for 359 diseases and injuries and healthy life expectancy (HALE) for 195 countries and territories, 1990-2017: a systematic analysis for the Global Burden of Disease Study 2017. Lancet. 2018;392(10159):1859–922.

7. Fischer A, Zimovetz E, Ling C, Esser D, and Schoof N. Humanistic and cost burden of systemic sclerosis: A review of the literature. Autoimmun Rev. 2017;16(11):1147–54.

8. Shapira Y, Agmon-Levin N, and Shoenfeld Y. Geoepidemiology of autoimmune rheumatic diseases. Nat Rev Rheumatol. 2010;6(8):468–76.

9. Denton CP, and Khanna D. Systemic sclerosis. Lancet. 2017;390(10103):1685–99.

10. Stoenoiu MS, Houssiau FA, and Lecouvet FE. Tendon friction rubs in systemic sclerosis: a possible explanation--an ultrasound and magnetic resonance imaging study. Rheumatology (Oxford). 2013;52(3):529–33.

11. Avouac J, Walker UA, Hachulla E, Riemekasten G, Cuomo G, Carreira PE, et al. Joint and tendon involvement predict disease progression in systemic sclerosis: a EUSTAR prospective study. Annals of the rheumatic diseases. 2014;75(1):103–9.

12. Dobrota R, Maurer B, Graf N, Jordan S, Mihai C, Kowal-Bielecka O, et al. Prediction of improvement in skin fibrosis in diffuse cutaneous systemic sclerosis: a EUSTAR analysis. Ann Rheum Dis. 2016;75(10):1743–8.

13. Distler JHW, Gyorfi AH, Ramanujam M, Whitfield ML, Konigshoff M, and Lafyatis R. Shared and distinct mechanisms of fibrosis. Nat Rev Rheumatol. 2019;15(12):705–30.

14. Galant C, Marchandise J, Stoenoiu MS, Ducreux J, De Groof A, Pirenne S, et al. Overexpression of ubiquitin-specific peptidase 15 in systemic sclerosis fibroblasts increases response to transforming growth factor beta. Rheumatology (Oxford). 2019;58(4):708–18.

15. Rooper LM, and Askin FB. In: Varga J, Denton CP, Wigley FM, Allanore Y, and Kuwana M eds. Scleroderma: From Pathogenesis to Comprehensive Management. Cham: Springer International Publishing; 2017:141–59.

16. Parker MW, Rossi D, Peterson M, Smith K, Sikström K, White ES, et al. Fibrotic extracellular matrix activates a profibrotic positive feedback loop. The Journal of clinical investigation. 2014;124(4):1622–35.

17. Bhattacharyya S, Tamaki Z, Wang W, Hinchcliff M, Hoover P, Getsios S, et al. FibronectinEDA promotes chronic cutaneous fibrosis through Toll-like receptor signaling. Science translational medicine. 2014;6(232).

18. Bhattacharyya S, Wang W, Morales-Nebreda L, Feng G, Wu M, Zhou X, et al. Tenascin-C drives persistence of organ fibrosis. Nat Commun. 2016;7:11703.

19. Herrera J, Henke CA, and Bitterman PB. Extracellular matrix as a driver of progressive fibrosis. The Journal of clinical investigation. 2018;128(1):45–53.

20. Tschumperlin DJ, Ligresti G, Hilscher MB, and Shah VH. Mechanosensing and fibrosis. J Clin Invest. 2018;128(1):74–84.

21. Lampi MC, and Reinhart-King CA. Targeting extracellular matrix stiffness to attenuate disease: From molecular mechanisms to clinical trials. Sci Transl Med. 2018;10(422):eaao0475.

22. Zitnay JL, Jung GS, Lin AH, Qin Z, Li Y, Yu SM, et al. Accumulation of collagen molecular unfolding is the mechanism of cyclic fatigue damage and failure in collagenous tissues. Science Advances. 2020;6(35).

23. Lavagnino M, and Arnoczky SP. In vitro alterations in cytoskeletal tensional homeostasis control gene expression in tendon cells. J Orthop Res. 2005;23(5):1211–8.

24. Gardner K, Lavagnino M, Egerbacher M, and Arnoczky SP. Re-establishment of cytoskeletal tensional homeostasis in lax tendons occurs through an actin-mediated cellular contraction of the extracellular matrix. Journal of Orthopaedic Research. 2012;30(11):1695–701.

25. Georges PC, Hui JJ, Gombos Z, McCormick ME, Wang AY, Uemura M, et al. Increased stiffness of the rat liver precedes matrix deposition: implications for fibrosis. American Journal of Physiology-Gastrointestinal and Liver Physiology. 2007;293(6):G1147–G54.

26. Wei SC, Fattet L, Tsai JH, Guo YR, Pai VH, Majeski HE, et al. Matrix stiffness drives epithelial mesenchymal transition and tumour metastasis through a TWIST1-G3BP2 mechanotransduction pathway. Nature Cell Biology. 2015;17(5):678–U306.

27. Vining KH, and Mooney DJ. Mechanical forces direct stem cell behaviour in development and regeneration. Nat Rev Mol Cell Biol. 2017;18(12):728–42.

28. Harris AK, Wild P, and Stopak D. Silicone rubber substrata: a new wrinkle in the study of cell locomotion. Science. 1980;208(4440):177–9.

29. Harris AK, Stopak D, and Wild P. Fibroblast traction as a mechanism for collagen morphogenesis. Nature. 1981;290(5803):249–51.

30. Sabass B, Gardel ML, Waterman CM, and Schwarz US. High resolution traction force microscopy based on experimental and computational advances. Biophys J. 2008;94(1):207–20.

31. Bergert M, Lendenmann T, Zundel M, Ehret AE, Panozzo D, Richner P, et al. Confocal reference free traction force microscopy. Nat Commun. 2016;7:12814.

32. Polacheck WJ, and Chen CS. Measuring cell-generated forces: a guide to the available tools. Nature methods. 2016;13(5):415–23.

33. Baker BM, and Chen CS. Deconstructing the third dimension - how 3D culture microenvironments alter cellular cues. Journal of Cell Science. 2012;125(13):3015–24.

34. Legant WR, Pathak A, Yang MT, Deshpande VS, McMeeking RM, and Chen CS. Microfabricated tissue gauges to measure and manipulate forces from 3D microtissues. Proc Natl Acad Sci U S A. 2009;106(25):10097–102.

35. Boudou T, Legant WR, Mu A, Borochin MA, Thavandiran N, Radisic M, et al. A microfabricated platform to measure and manipulate the mechanics of engineered cardiac microtissues. Tissue Eng Part A. 2012;18(9-10):910–9.

36. Sakar MS, Neal D, Boudou T, Borochin MA, Li Y, Weiss R, et al. Formation and optogenetic control of engineered 3D skeletal muscle bioactuators. Lab Chip. 2012;12(23):4976–85.

37. Sakar MS, Eyckmans J, Pieters R, Eberli D, Nelson BJ, and Chen CS. Cellular forces and matrix assembly coordinate fibrous tissue repair. Nat Commun. 2016;7:11036.

38. Eyckmans J, and Chen CS. 3D culture models of tissues under tension. J Cell Sci. 2017;130(1):63–70.

39. Davidson MD, Prendergast ME, Ban E, Xu KL, Mickel G, Mensah P, et al. Programmable and Contractile Materials Through Cell Encapsulation in Fibrous Hydrogel Assemblies. bioRxiv. 2021:2021.04.19.440470.

40. Schoen I, Pruitt BL, and Vogel V. The Yin-Yang of Rigidity Sensing: How Forces and Mechanical Properties Regulate the Cellular Response to Materials. Annual Review of Materials Research, Vol 43. 2013;43:589–618.

41. Kalson NS, Lu Y, Taylor SH, Starborg T, Holmes DF, and Kadler KE. A structure-based extracellular matrix expansion mechanism of fibrous tissue growth. Elife. 2015;4.

42. Subramanian A, Kanzaki LF, Galloway JL, and Schilling TF. Mechanical force regulates tendon extracellular matrix organization and tenocyte morphogenesis through TGFbeta signaling. Elife. 2018;7.

43. Polacheck WJ, and Chen CS. Measuring cell-generated forces: a guide to the available tools. Nat Methods. 2016;13(5):415–23.

44. Yang C, Tibbitt MW, Basta L, and Anseth KS. Mechanical memory and dosing influence stem cell fate. Nat Mater. 2014;13(6):645–52.

45. Li CX, Talele NP, Boo S, Koehler A, Knee-Walden E, Balestrini JL, et al. MicroRNA-21 preserves the fibrotic mechanical memory of mesenchymal stem cells. Nat Mater. 2017;16(3):379–89.

46. Whitfield ML, Finlay DR, Murray JI, Troyanskaya OG, Chi JT, Pergamenschikov A, et al. Systemic and cell type-specific gene expression patterns in scleroderma skin. Proc Natl Acad Sci U S A. 2003;100(21):12319–24.

47. Sargent JL, Milano A, Bhattacharyya S, Varga J, Connolly MK, Chang HY, et al. A TGFbeta-responsive gene signature is associated with a subset of diffuse scleroderma with increased disease severity. J Invest Dermatol. 2010;130(3):694–705.

48. Palumbo-Zerr K, Zerr P, Distler A, Fliehr J, Mancuso R, Huang J, et al. Orphan nuclear receptor NR4A1 regulates transforming growth factor-beta signaling and fibrosis. Nat Med. 2015;21(2):150–8.

49. Lofgren S, Hinchcliff M, Carns M, Wood T, Aren K, Arroyo E, et al. Integrated, multicohort analysis of systemic sclerosis identifies robust transcriptional signature of disease severity. Jci Insight. 2016;1(21).

50. Bhattacharyya S, Wang W, Qin W, Cheng K, Coulup S, Chavez S, et al. TLR4-dependent fibroblast activation drives persistent organ fibrosis in skin and lung. JCI Insight. 2018;3(13):e98850.

51. Shin JY, Beckett JD, Bagirzadeh R, Creamer TJ, Shah AA, McMahan Z, et al. Epigenetic activation and memory at a TGFB2 enhancer in systemic sclerosis. Science Translational Medicine. 2019;11(497).

52. Goodier HC, Carr AJ, Snelling SJ, Roche L, Wheway K, Watkins B, et al. Comparison of transforming growth factor beta expression in healthy and diseased human tendon. Arthritis Res Ther. 2016;18:48.

53. Morita W, Snelling SJ, Dakin SG, and Carr AJ. Profibrotic mediators in tendon disease: a systematic review. Arthritis Res Ther. 2016;18(1):269.

54. Wang X, Xie L, Crane J, Zhen G, Li F, Yang P, et al. Aberrant TGF-beta activation in bone tendon insertion induces enthesopathy-like disease. J Clin Invest. 2018;128(2):846–60.

55. Gabbiani G, Ryan GB, and Majne G. Presence of modified fibroblasts in granulation tissue and their possible role in wound contraction. Experientia. 1971;27(5):549–50.

56. Hinz B, Celetta G, Tomasek JJ, Gabbiani G, and Chaponnier C. Alpha-smooth muscle actin expression upregulates fibroblast contractile activity. Mol Biol Cell. 2001;12(9):2730–41.

57. Tomasek JJ, Gabbiani G, Hinz B, Chaponnier C, and Brown RA. Myofibroblasts and mechano-regulation of connective tissue remodelling. Nat Rev Mol Cell Biol. 2002;3(5):349–63.

58. Hinz B, and Lagares D. Evasion of apoptosis by myofibroblasts: a hallmark of fibrotic diseases. Nature Reviews Rheumatology. 2020;16(1):11–31.

59. Parker MW, Rossi D, Peterson M, Smith K, Sikstrom K, White ES, et al. Fibrotic extracellular matrix activates a profibrotic positive feedback loop. J Clin Invest. 2014;124(4):1622–35.

60. Avouac J, Walker UA, Hachulla E, Riemekasten G, Cuomo G, Carreira PE, et al. Joint and tendon involvement predict disease progression in systemic sclerosis: a EUSTAR prospective study. Ann Rheum Dis. 2016;75(1):103–9.

61. Eferl R, Hasselblatt P, Rath M, Popper H, Zenz R, Komnenovic V, et al. Development of pulmonary fibrosis through a pathway involving the transcription factor Fra-2/AP-1. Proceedings of the National Academy of Sciences. 2008;105(30):10525–30.

62. Maurer B, Reich N, Juengel A, Kriegsmann J, Gay RE, Schett G, et al. Fra-2 transgenic mice as a novel model of pulmonary hypertension associated with systemic sclerosis. Annals of the Rheumatic Diseases. 2012;71(8):1382–7.

63. Ucero AC, Bakiri L, Roediger B, Suzuki M, Jimenez M, Mandal P, et al. Fra-2-expressing macrophages promote lung fibrosis in mice. J Clin Invest. 2019;129(8):3293–309.

64. Beyer C, Schett G, Distler O, and Distler JH. Animal models of systemic sclerosis: prospects and limitations. Arthritis Rheum. 2010;62(10):2831–44.

65. Renoux F, Stellato M, Impellizzieri D, Huang R, Subramaniam A, Dees C, et al. Arthritis Rheumatol. 2016.

66. Wunderli SL, Widmer J, Amrein N, Foolen J, Silvan U, Leupin O, et al. Minimal mechanical load and tissue culture conditions preserve native cell phenotype and morphology in tendon-a novel ex vivo mouse explant model. J Orthop Res. 2018;36(5):1383–90.

67. Renoux F, Stellato M, Haftmann C, Vogetseder A, Huang RY, Subramaniam A, et al. The AP1 Transcription Factor Fosl2 Promotes Systemic Autoimmunity and Inflammation by Repressing Treg Development. Cell Reports. 2020;31(13).

68. Goffin JM, Pittet P, Csucs G, Lussi JW, Meister JJ, and Hinz B. Focal adhesion size controls tension-dependent recruitment of alpha-smooth muscle actin to stress fibers. J Cell Biol. 2006;172(2):259–68.

69. Cukierman E, Pankov R, Stevens DR, and Yamada KM. Taking cell-matrix adhesions to the third dimension. Science. 2001;294(5547):1708–12.

70. Wipff PJ, Rifkin DB, Meister JJ, and Hinz B. Myofibroblast contraction activates latent TGF-beta1 from the extracellular matrix. J Cell Biol. 2007;179(6):1311–23.

71. Wu HJ, Yu YY, Huang HW, Hu YC, Fu SL, Wang Z, et al. Progressive Pulmonary Fibrosis Is Caused by Elevated Mechanical Tension on Alveolar Stem Cells. Cell. 2020;180(1):107-+.

72. Liu F, Mih JD, Shea BS, Kho AT, Sharif AS, Tager AM, et al. Feedback amplification of fibrosis through matrix stiffening and COX-2 suppression. Journal of Cell Biology. 2010;190(4):693–706.

73. Sharma RI, and Snedeker JG. Biochemical and biomechanical gradients for directed bone marrow stromal cell differentiation toward tendon and bone. Biomaterials. 2010;31(30):7695–704.

74. Podolsky MJ, Yang CD, Valenzuela CL, Datta R, Huang SK, Nishimura SL, et al. Age-dependent regulation of cell-mediated collagen turnover. Jci Insight. 2020;5(10).

75. Angelidis I, Simon LM, Fernandez IE, Strunz M, Mayr CH, Greiffo FR, et al. An atlas of the aging lung mapped by single cell transcriptomics and deep tissue proteomics. Nature Communications. 2019;10.

76. Sicard D, Haak AJ, Choi KM, Craig AR, Fredenburgh LE, and Tschumperlin DJ. Aging and anatomical variations in lung tissue stiffness. American Journal of Physiology-Lung Cellular and Molecular Physiology. 2018;314(6):L946–L55.

77. Segel M, Neumann B, Hill MFE, Weber IP, Viscomi C, Zhao C, et al. Niche stiffness underlies the ageing of central nervous system progenitor cells. Nature. 2019;573(7772):130-+.

78. Beyeler F, Neild A, Oberti S, Bell DJ, Sun Y, Dual J, et al. Monolithically Fabricated Microgripper With Integrated Force Sensor for Manipulating Microobjects and Biological Cells Aligned in an Ultrasonic Field. Journal of Microelectromechanical Systems. 2007;16(1).

79. Klotzsch E, Smith ML, Kubow KE, Muntwyler S, Little WC, Beyeler F, et al. Fibronectin forms the most extensible biological fibers displaying switchable force-exposed cryptic binding sites. Proceedings of the National Academy of Sciences of the United States of America. 2009;106(43):18267–72.

80. Poulsen RC, Carr AJ, and Hulley PA. Protection against Glucocorticoid-Induced Damage in Human Tenocytes by Modulation of ERK, Akt, and Forkhead Signaling. Endocrinology. 2011;152(2):503–14.

81. Phelan K, and May KM. Basic Techniques in Mammalian Cell Tissue Culture. Current Protocols in Cell Biology. 2015;66(1):1. 22.

82. Murchison ND, Price BA, Conner DA, Keene DR, Olson EN, Tabin CJ, et al. Regulation of tendon differentiation by scleraxis distinguishes force-transmitting tendons from muscle-anchoring tendons. Development (Cambridge, England). 2007;134(14):2697–708.

83. Rajan N, Habermehl J, Cote MF, Doillon CJ, and Mantovani D. Preparation of ready-to-use, storable and reconstituted type I collagen from rat tail tendon for tissue engineering applications. Nat Protoc. 2006;1(6):2753–8.

84. Weischenfeldt J, and Porse B. Bone Marrow-Derived Macrophages (BMM): Isolation and Applications. Cold Spring Harbor Protocols. 2008;2008(12):pdb.prot5080.

85. Schmittgen TD, and Livak KJ. Analyzing real-time PCR data by the comparative C(T) method. Nat Protoc. 2008;3(6):1101–8.

